# Dynamic Mechanism for Subtype Selectivity of Endocannabinoids

**DOI:** 10.1101/2024.10.25.620304

**Authors:** Soumajit Dutta, Lawrence Zhao, Diwakar Shukla

## Abstract

Endocannabinoids are naturally occurring lipid-like molecules that bind to cannabinoid receptors (CB_1_ and CB_2_) and regulate many of human bodily functions via the endocannabinoid system. There is a tremendous interest in developing selective drugs that target the CB receptors. However, the biophysical mechanisms responsible for the subtype selectivity for endocannbinoids have not been established. Recent experimental structures of CB receptors show that endocannbinoids potentially bind via membrane using the lipid access channel in the transmembrane region of the receptors. Furthermore, the N-terminus of the receptor could move in and out of the binding pocket thereby modulating both the pocket volume and its residue composition. On the basis of these observations, we propose two hypothesis to explain the selectivity of the endocannabinoid, anandamide for CB_1_ receptor. First, the selectivity arises from distinct enthalpic ligand-protein interactions along the ligand binding pathway formed due to the movement of N-terminus and subsequent shifts in the binding pocket composition. Second, selectivity arises from the volumetric differences in the binding pocket allowing for differences in ligand conformational entropy. To quantitatively test these hypothesis, we perform extensive molecular dynamics simulations (∼0.9 milliseconds) along with Markov state modeling and deep learning-based VAMPnets to provide an interpretable characterization of the anandamide binding process to cannabinoid receptors and explain its selectivity for CB_1_. Our findings reveal that the distinct N-terminus positions along lipid access channels between TM1 and TM7 lead to different binding mechanisms and interactions between anandamide and the binding pocket residues. To validate the critical stabilizing interactions along the binding pathway, relative free energy calculations of anandamide analogs are used. Moreover, the larger CB_2_ pocket volume increases the entropic effects of ligand binding by allowing higher ligand fluctuations but reduced stable interactions. Therefore, the opposing enthalpy and entropy effects between the receptors shape the endocannabinoid selectivity. Overall, the CB_1_ selectivity of anandamide is explained by the dominant enthalpy contributions due to ligand-protein interactions in stable binding poses. This study shed lights on potential selectivity mechanisms for endocannabinoids that would aid in the discovery of CB selective drugs.

## Introduction

The discovery of cannabinoid receptors (CBRs) busted the myth of “membrane-mediated signaling” of lipid-like cannabinoid molecules.^1^ Two types of cannabinoid receptors have been confirmed: Cannabinoid receptor 1 (CB_1_) and 2 (CB_2_).^2^ CB_1_ is majorly expressed in central nervous systems, whereas CB_2_ is primarily expressed in the immune system.^3,4^ The discoveries of CBRs established a complex signaling system named the endocannabinoid system (ECS).^5,6^ The ECS consists of neurotransmitter endocannabinoids, enzymes that help produce or degrade endocannabinoids, and receptors where the endocannabinoids bind to modulate downstream signaling.^7^ This complex system maintains homeostasis in neuronal signaling and has been targeted for different physiological and psychological disorders such as obesity, pain, and inflammation.^8^

Unlike other neurotransmitters, which are stored for future usage, endocannabinoids are produced in demand from the membranes of postsynaptic neurons due to an increase in calcium ion concentration.^9^ These molecules travel to the presynaptic neuron via a retrograde signaling system to bind CBRs.^10^ Binding of endocannabinoid leads to downstream signaling by G-protein and *β*-arrestin.^11–13^ Enzymes that are part of ECS finally degrade these molecules in presynaptic neurons. ^14^

Two well-known endocannabinoids are anandamide (N-Arachidonoylethanolamine) and 2-AG (2-arachidonoylglycerol) (Figure 1A). ^17,18^ These molecules have four pharmacophore groups: a polar head group, a propyl linker, a polyene linker, and an alkyl tail. Although both molecules are precursors of arachidonic acid, they are produced via distinct enzymatic mechanisms.^19,20^ Anandamide is a partial agonist for both CB_1_ and CB_2_, whereas 2-AG is a full agonist for both receptors.^21^ As full agonists of CBRs are associated with higher side effects, anandamide has gained more attention in drug design due to its partial agonistic properties like Δ^9^-THC.^22–24^

**Figure 1:**
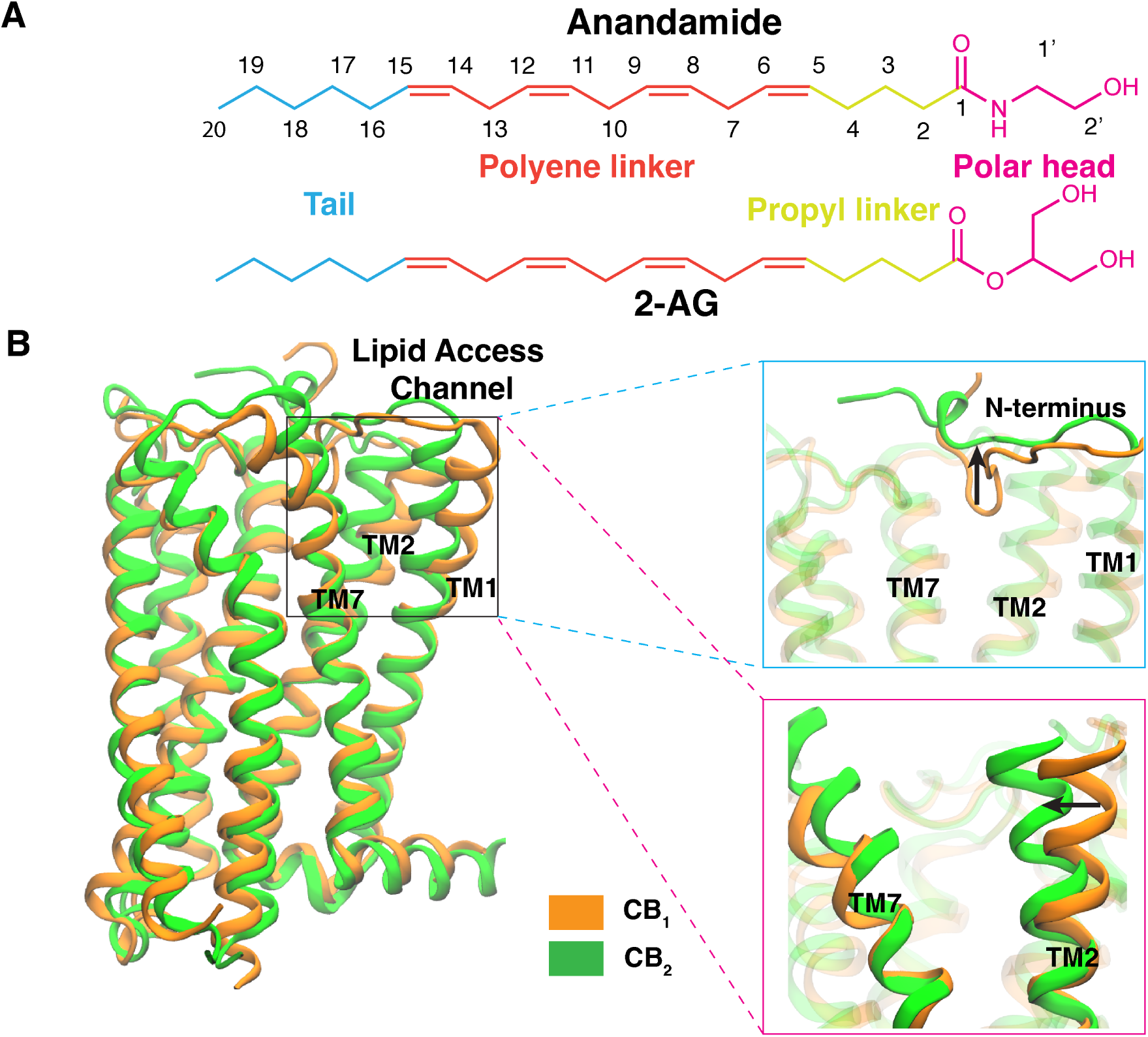
(A) Structures of the two well-known endocannabinoids: anandamide and 2-AG. Four pharmacophore groups of each endocannabinoid are represented in different colors. The carbon atom numbering of anandamide is shown. (B) Superposition of inactive CB_1_ (PDB: 5TGZ; ^15^ color: orange) and CB_2_ (PDB: 5ZTY; ^16^ color: green) are shown as cartoon representations with structural differences along the lipid access channel highlighted in separate panels.

GPCRs belonging to the same subfamily share high sequence and structural similarities.^25^ Consequently, most designed drugs bind to all receptors from the same subfamily, causing off-target side effects. CB_1_ and CB_2_ share a 44% total sequence identity and 68% sequence homology in the transmembrane regions.^26^ Previous experiments have shown anandamide to be selective towards CB_1_ (*K_i_*: 61-543 nM) as compared to CB_2_ (*K_i_*: 279-1940 nM).^18,21^ Several anandamide analogs designed to study the structure-activity relationship have been shown to retain or increase ligand selectivity for CB_1_.^27–29^ Therefore, anandamide and its analogs provide an avenue to design subtype-selective ligands for CB_1_, which is an important issue not just for CBRs but for all Class A GPCRs.^30–32^

This study proposes two hypotheses to provide mechanistic explanation of endocannabinoid subtype selectivity. The first hypothesis focuses on the influence of structural changes along the endocannabinoid binding pathway within the receptors. The second hypothesis explores the role of entropy in the binding process.

Previous studies suggested that endocannabinoids, due to their lipidic nature, bind to the CBRs from the membrane-accessible transmembrane regions via lateral diffusion.^33,34^ This proposition was supported by the recent discoveries of experimentally determined structures that show a lipid accession channel between TM1 and TM7 for CB_1_.^22,35^ Additionally, mutagenesis and molecular dynamics (MD) studies on cannabinoid receptors provide further evidence for this ligand binding pathway.^33,36,37^ However, comparison of the lipid access channel of CB_1_ and CB_2_ inactive structures shows major structural differences (Figure 1B). Membrane-proximal N-terminus remains inside the orthosteric pocket for CB_1_ inactive structure.^15,38^ Agonist binding moves the N-terminus out of the pocket in order to bind in the agonist-like pose.^23,35^ Conversely, the downward N-terminus creates a large space between TM2 and TM7 for CB_1_ compared to CB_2_, which might facilitate the entrance of the agonists in the pocket (Figure 1B). These structural differences might influence endocannabinoid binding mechanism and ligand interactions in bound pose within each receptor. Thus, in our first hypothesis, we propose the distinct interactions formed by the ligand and receptors lead to differential enthalpy contributions to binding affinity making anandamide subtype selective.

The second hypothesis is based on the volumetric differences in cannabinoid receptors’ binding pocket.^38^ CB_1_ and CB_2_ binding pockets have been shown to accommodate much bulkier ligands compared to the long-chain endocannabinoids with an abundance of *sp*^3^ hybridized carbon atoms.^15,35^ Flexible endocannabinoids may obtain multiple reversible stable poses inside large pocket of CBRs, which increases the importance of entropy contribution in ligand binding. Recent experimental studies have also revealed that entropy plays a major role in determining subtype selectivity of CBRs.^39^ Therefore, we propose that distinct pocket volumes of CBRs leads to dissimilar entropy contribution in binding free energy, causing subtype selective behavior of anandamide. Hence, to investigate whether distinct ligand-protein interactions or ligand flexibility effects determine ligand selectivity, we compared the binding of the endocannabinoid anandamide to cannabinoids using atomistic MD simulations.

Extensive ∼ 900*µ*s of unbiased molecular dynamics were performed for the exploration of binding pathways and interaction sites of anandamide for CB_1_ and CB_2_. The thermodynamics and kinetics of these processes were estimated using Markov state models (MSM). ^40^ Deep learning based VAMPnets was used to identify intermediate states along the pathway.^41^ Binding simulations reveal that anandamide can diffuse through the membrane to bind to CB_1_ and CB_2_ from the lipid access channel of TM1 and TM7 confirming previous observations. A significant conformational change in membrane-proximal N-terminus of CB_1_ during ligand binding suggests an induced fit binding mechanism. Conversely, anandamide binding to CB_2_ does not lead to a major conformational change in the binding pocket, aligning with a conformational selection mechanism. Unbiased MD simulations also detect a non-canonical transmembrane pathway only for CB_2_ (between TM5 and TM6), which converges to the orthosteric binding pocket similar to the canonical pathway but appears less favorable for ligand binding. This difference in binding mechanisms contributes to varying interactions within the binding pockets of CB_1_ and CB_2_. These interaction differences were validated by relative binding free energy (RBFE) calculations for anandamide analogs, which show consistent trends with experimental binding affinity. Further analyses reveal that anandamide interacts strongly with the displaced N-terminus in the orthosteric pocket of CB_1_, increasing enthalpy contribution to binding free energy compared to CB_2_, where the N-terminus remains outside the pocket. Additionally, larger volume of orthosteric binding pocket of CB_2_ allow more flexibility for the anandamide compared to CB_1_ in the ligand bound state. This increased flexibility leads to a higher entropy contribution towards ligand binding free energy for CB_2_. Comparing these opposing entropy and enthalpy contributions reveals that the enthalpic contributions are dominant and facilitate more selective ligand binding in CB_1_. Overall, this study provides a detailed quantitative evaluation of mechanistic hypothesis for understanding of endocannabinoid subtype selectivity, which will aid in the discovery of selective drugs targeting cannabinoid receptors.

## Results and Discussion

### Anandamide binds to cannabinoid receptors via the lipid access channels in between transmembrane helices

Approximately ∼ 423 and 483 *µ*s of atomistic molecular dynamics simulations were performed for CB_1_ and CB_2_ to capture the anandamide binding. Realistic membrane compositions were used to accurately estimate thermodynamics and kinetics as the ligand binding happens via the membrane bilayer. ^42,43^ Average brain membrane composition was used for CB_1_ as these receptors are majorly expressed in the central nervous system (Table S1). In contrast, average eukaryotic membrane composition was used for CB_2_ as they are expressed in immune cells (Table S2). Anandamide was initially placed in the extracellular region and was observed to diffuse to the membrane bilayer for both CB_1_ and CB_2_ during the simulation. The higher probability density of the ligand in the membrane bilayer compared to the salt solution is consistent with the lipidic nature of anandamide (Figure S1). The frames where ligand stays in the membrane bilayer were selected to characterize its transmembrane diffusion and binding pathway to the receptors. Markov State Models were used to obtain the unbiased free energy estimates of the ligand binding process. MSM-weighted free energy landscape between *x* and *y* components of the center of mass of anandamide reveal a binding pathway for cannabinoid receptors (CB_1_ and CB_2_) from the lipid access channel between TM1 and TM7 (Figure 2A and 2B). A separate binding pathway was also observed in between TM5 and TM6 for CB_2_, which converges to the similar bound pose inside the pocket (Figure 2B).

**Figure 2:**
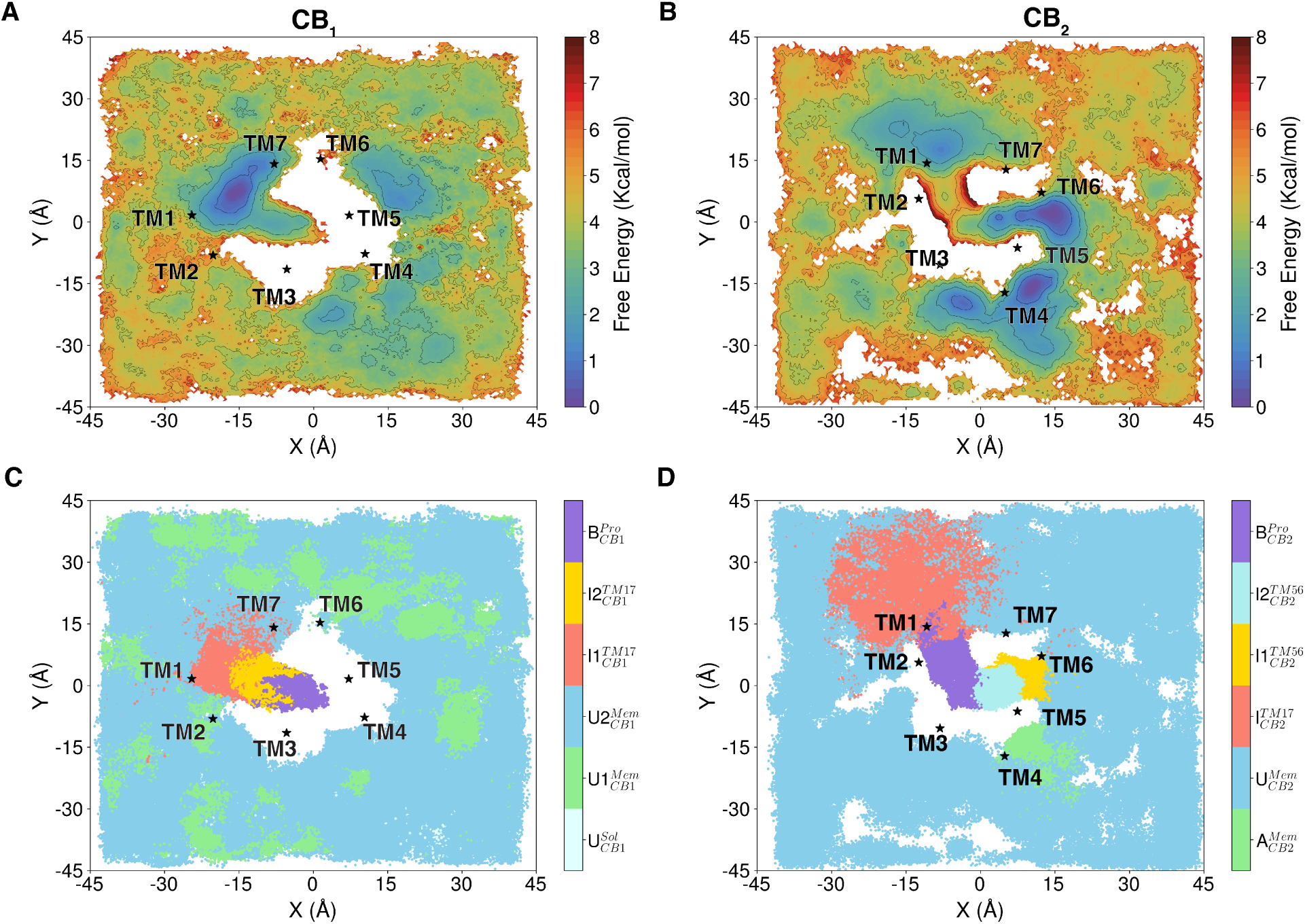
MSM weighted free energy landscapes are projected between *x* and *y* coordinates of center of mass of anandamide for CB_1_ (A) and CB_2_ (B). Macrostates obtained by training the VAMPnets are plotted as scatter plots between *x* and *y* coordinates of the center of mass of anandamide for CB_1_ (C) and CB_2_ (D). Frames from the transmembrane region are only shown in the above plots. The extracellular residue of each helix is shown as a star marker to show its position. Macrostate names A, U, I, and B represent anandmide in allosterically bound, unbound, intermediate and orthosterically bound state in the receptors. Superscript in the intermediate macrostate stands for the ligand binding pathway between transmembrane helices.

Characterization of the binding process was done using VAMPnets, which cluster the simulation ensemble into different macrostates. VAMPnets use deep learning architecture to divide the ensemble into macrostates, where intrastate conformational changes are faster compared to interstate transitions.^41^ The optimal number of macrostates were selected for CB_1_ (six) and CB_2_ (six) with the criteria of minimizing the error in implied timescale and state populations (Figures S2, S3, S4, and S5). Relative positions of the macrostates are shown as scatter plots based on the *x* and *y* components of the ligand center of mass of frames where anandamide stay inside membrane-bilayer or protein (Figures 2C and 2D).

The agonist bound pose in CB_1_ and CB_2_ are characterized by the macrostate *B^Pro^_CB1_* and *B^Pro^_CB2_*, respectively. Two intermediate states (*I*1*^TM17^_CB1_*, *I*2*^TM17^_CB1_*) mark the anandamide binding pathway in between TM1 and TM7 for CB_1_ before the ligand reach bound state from the bulk membrane. In contrast, one intermediate state (*I^TM^*^17^_CB2_) was observed along the similar pathway for CB_2_. An alternate pathway was also observed for CB_2_ in between TM5 and TM6 was characterized by two intermediate states (*I*1*^TM56^_CB2_*, *I*2*^TM56^_CB2_*). Detailed description of each pathway and its role in determining selectivity are explained in the subsequent sections.

### Induced-fit binding of anandamide to CB_1_ via the lipid access channel of TM1 and TM7

Ligand diffuse from the bulk membrane to the metastable minima at the membrane exposed interface of TM1 and TM7 before anandamide binds inside the pocket (Figure 2A). This stable region of the landscape is represented as Macrostate *I*1*^TM17^_CB1_* (Figure 2C). The timescale for ligand diffusion from the bulk of the membrane was determined using mean free passage time (MFPT) calculation. MFPT was calculated from transition path theory, which estimates the timescale from the transition probability matrix as determined by MSM. MFPT calculation shows that the ligand can diffuse from bulk to microstate *I*1*^TM17^_CB1_* within a sub-microsecond timescale (0.6 ± 0.1*µ*s).

In this metastable macrostate *I*1*^TM17^_CB1_*, the ligand is stabilized in the protein-membrane surface while maintaining contact with the lipid molecules (Figures 3A and 3B). Protein and lipid exposed surface area calculation of anandamide in macrostate *I*1*^TM17^_CB1_* shows that more than 50% of average area maintains contact with protein, depicting a stable interaction between the ligand and protein (Figure S6). The protein-ligand contact probabilities and interaction energies were calculated to identify the residues important for anandamide stabilization at the receptor surface (Figure 4A). These calculations reveal that the ligand form major interactions with N-terminus (F108*^N^*^−*term*^, M109*^N^*^−*term*^) and TM7 (F381^7.37^, M384^7.40^) residues (Figure 3C). The most frequent and stable interaction is the polar hydrogen bond interaction between the nitrogen group of anandamide and backbone oxygen of the N-terminus F108*^N^*^−*term*^ residue. Contact probability calculations were also performed to capture major interactions of anandamide pharmacophore groups with different lipids (Figure 3B). The lipids in the membrane were classified based on the eight different head groups. The analysis results reveal that major interactions with the ligand are observed with phosphatidylcholine (PC), sphingolipids (SM), and cholesterol (CHL). Furthermore, stable lipid contacts are majorly formed with the polyene linker and polar headgroups, while the propyl chain and hydrophobic tail form weaker interactions (Figure 3B). Due to the stable interactions, the probability density plot for ligand RMSD reveals only one major conformation of anandamide, where the ligand is parallel to the protein helices (Figures 3C and S7).

**Figure 3:**
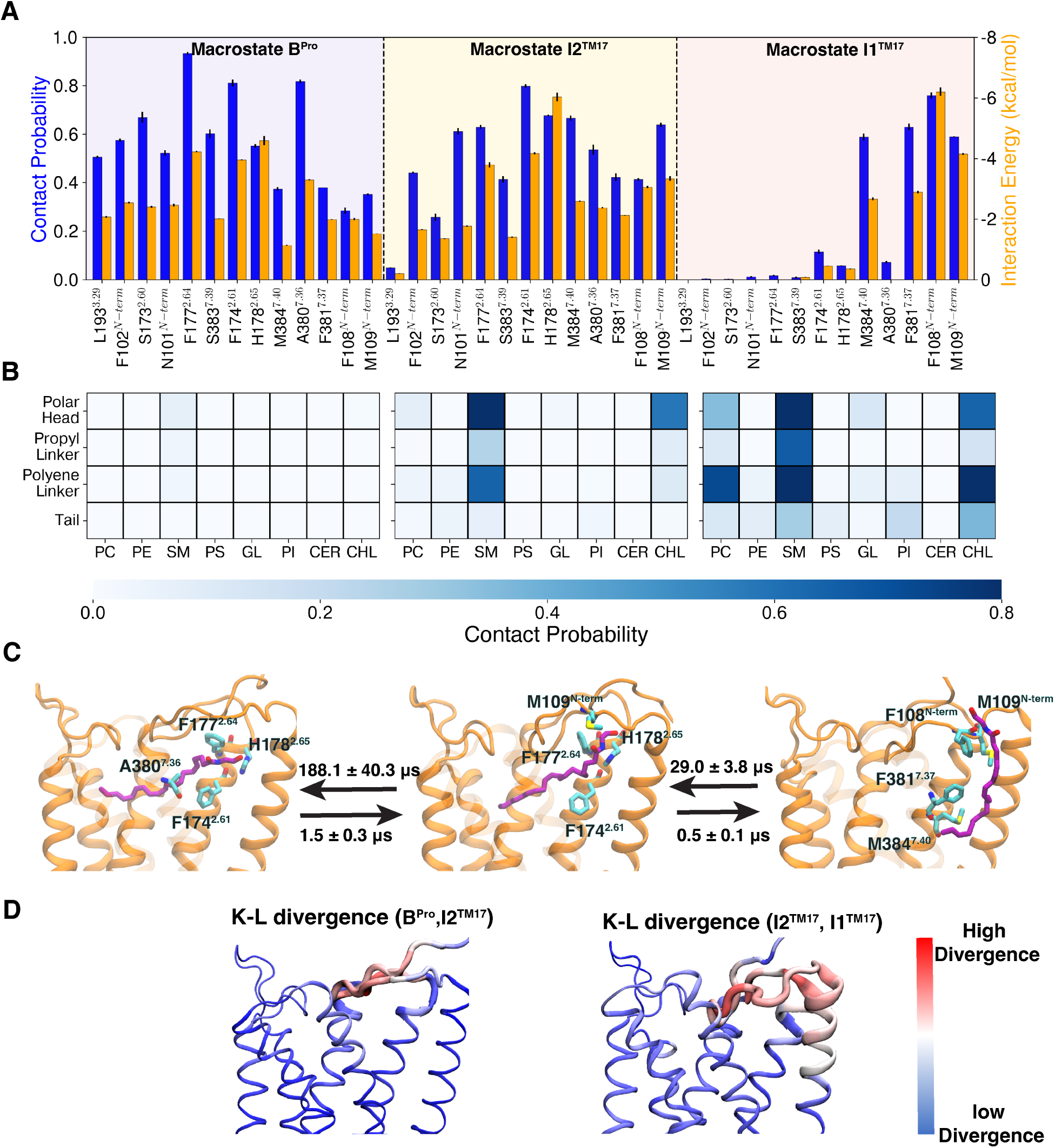
(A) Contact probability (color: Blue) and interaction energy (color: Orange) calculations between anandamide and CB_1_ in different macrostates in the binding pathway. Residues that maintain at least 50% contact probabilities in one of the macrostates are shown in the figure. (B) Lipid contact probabilities in different macrostates of CB_1_ with anandamide are shown as a heatmap. (C) A representative frame from each of the macrostates is shown. CB_1_ (color: Orange) is shown as a cartoon representation with transparent TM6 and TM7. Anandamide (color: Purple) and the four highest interacting residues (color: Cyan) are represented as sticks. (D) K-L divergences between two macrostates are shown in color and thickness gradients. Thickness gradients are shown as moving averages.

**Figure 4:**
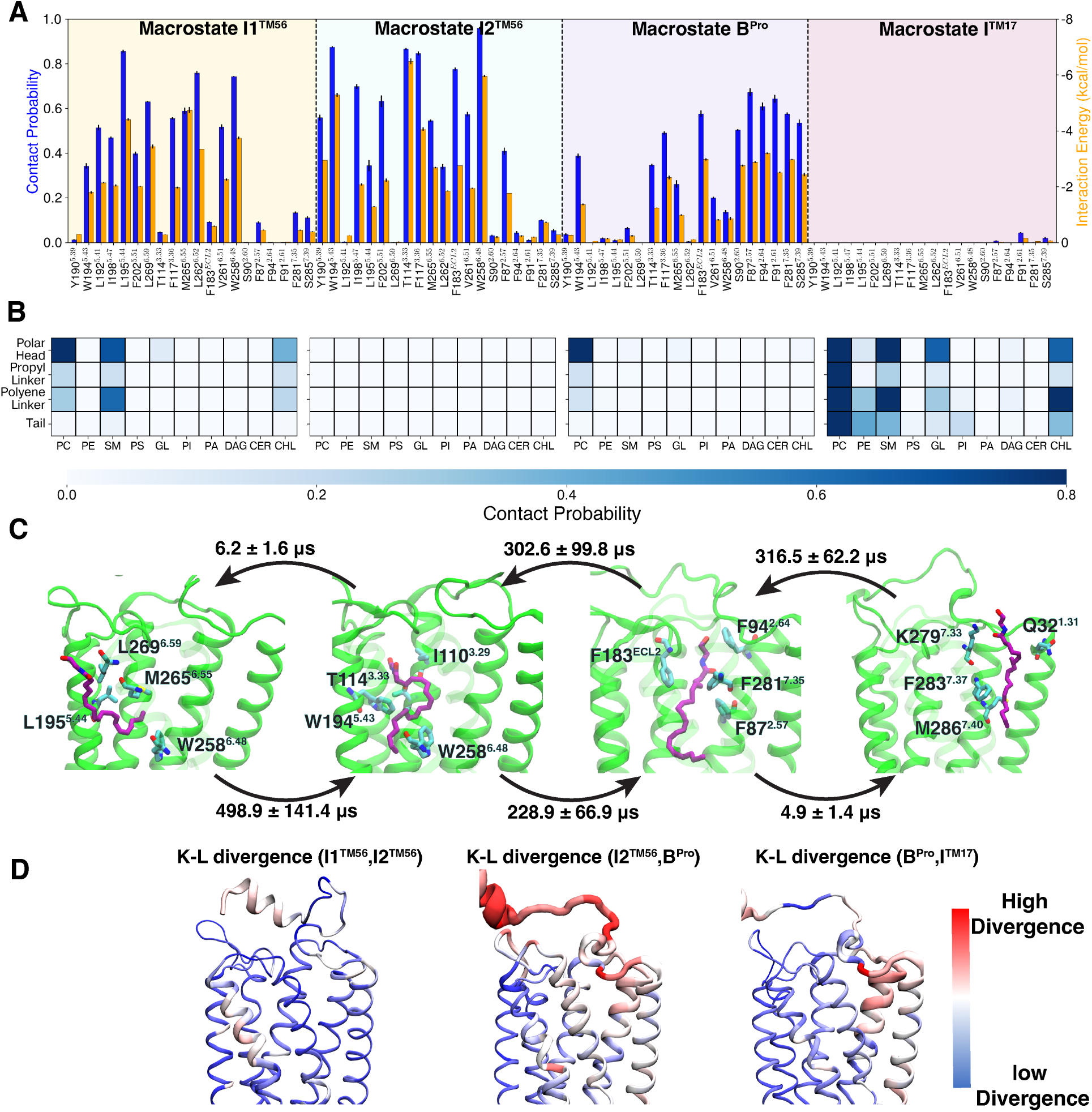
(A) Contact probability (color: Blue) and interaction energy (color: Orange) calculations between anandamide and CB_1_ in different macrostates in the binding pathway. Residues which maintain atleast 50% contact probabilities in one of the macrostates are shown in the figure. (B) Lipid contact probabilities in different macrostates of CB_1_ with anandamide are shown as a heatmap. (C) Representative frame from the each of the macrostate is shown. CB_1_ (color: Green) are shown as a cartoon representation with transparent TM6 and TM7. Anandamide (color: Purple) and four highest interacting residues (color: Cyan) are represented as sticks. (D) K-L divergences between two macrostates are shown color and thickness gradients. Thickness gradients are shown as moving average.

From macrostate *I*1*^TM17^_CB1_*, anandamide moves inside the binding pocket between the space formed by TM1, TM2, and TM7. Conformational ensemble of anandamide in this position was represented by macrostate *I*2*^TM17^_CB1_*. The binding position of anandamide in this macrostate is similar to the experimentally determined antagonist-bound pose (Figure S8). Contact probability and interaction energy calculations reveal that major ligand-protein interactions are formed between the polar group of the ligand with TM2 (F174^2.61^, F177^2.64^, and H178^2.65^) and N-terminus (M109*^N^*^−*term*^) of the protein (Figures 3A and 3C). Previous experiments have shown that these interacting residues in TM2 have major effects on the ligand binding and potency for CB_1_.^15,44^ Few lipid interactions are also present in the macrostate *I*2*^TM17^_CB1_* (Figure 3B). The polar part of the ligand still maintains contact with cholesterol (CHL) and sphingolipids (SM). Although macrostate *I*2*^TM17^_CB1_* has more protein interactions than macrostate *I*1*^TM17^_CB1_*, it was observed from the free energy landscape that macrostate *I*2*^TM17^_CB1_* is comparatively less stable than macrostate *I*1*^TM17^_CB1_* (Figure 2A). This observation shows the importance of lipid interactions in stabilizing anandamide in the protein interface. As the ligand moves from a low to high energy macrostate, kinetically, this transition is slower (29.0 ± 3.8*µ*s) compared to the reverse transition (0.5 ± 0.1*µ*s).

The binding of the ligand to the protein leads to the conformational change. K-L divergence analysis was performed between the different macrostates to highlight the regions of the protein conformational differences. Comparing the K-L divergence between these macrostates, a major conformational change was observed in the extracellular TM1, TM2, and N-terminus, confirming that the ligand binding is leading local conformational changes in the binding pocket (Figure 3D). Within the local conformational changes, specific movements are found to facilitate the ligand binding: upward movement of N-terminus and F174^2.61^. To capture these changes, the N-terminus (M103*^N^*^−*term*^) distance from D163^2.50^ was plotted against the sidechain dihedral (*χ*_1_) of F174^2.61^ as MSM weighted free energy landscape (Figures S9A and S9B). We observed that macrostate *I*1*^TM17^_CB1_* to macrostate *I*2*^TM17^_CB1_* transition leads to the shift in the population distribution of the *χ*_1_ angle of F174^2.61^. The binding of the macrostate *I*2*^TM17^_CB1_* also partially shifts the N-terminus in the extracellular direction to accommodate the ligand (Figure S9A).

From macrostate *I*2*^TM17^_CB1_*, anandamide moves further deep in the pocket to macrostate *B^Pro^_CB1_*, which represents the agonist binding pocket (Figure 3C). The major interactions are still maintained between the polar group of anandamide and TM2 in this macrostate (Figure 3A). K-L divergence calculation between these macrostates also shows the major conformational change associated with the N-terminus (Figure 3D). MSM weighted free energy landscape also points out the population shift of N-terminus away from the binding pocket (Figures S9A and S9B). This required large conformational change of the N-terminus in the binding pocket makes transition from *I*2*^TM17^_CB1_* order of magnitude slower compared to previous transitions along this pathway.

These significant conformational changes in the N-terminus alter the shape of the binding pocket. Further analyses reveal changes in the pocket volume during ligand binding (Figure S10). These observations suggest that anandamide binds to CB_1_ via an induced fit mechanism, where the binding of anandamide induces a conformational change in the N-terminus.

### Conformational selection mechanism dominates anandamide binding to CB_2_ via TM1-TM7 lipid access channel

Mutagenesis experiment and MD simulation have also shown similar ligand binding pathway in between TM1 and TM7 for CB_2_. However, the N-terminus of CB_2_ remain above the orthosteric pocket and far from this pathway.

Simulations show this structural distinction causes major difference in the pathway. VAMPnets analysis captures one intermediate macrostate *I^TM^*^17^ near the protein surface between TM1 and TM7 before anandamide binds to agonist-like poses in the orthosteric pocket. Spontaneous transition to this macrostate from membrane-bilayer was observed within the microsecond timescale (8.7 ± 1.2*µ*s). Contrasting to CB_1_, this macrostate does not represent a metastable minima in the protein surface between TM1 and TM7, which depicts less stable interactions with the protein surface (Figure 2B). The less stability of the ligand in this receptor surface can be explained by the lack of stable contact in the surface (Figures 4A). A lesser ratio of protein and lipid accessible surface area of anandamide compared to the Macrostate *I*1*^TM17^_CB1_* further validates this phenomena (Figure S6). Major interaction differences between CB_1_ and CB_2_ for anandamide was observed at the N-terminus. Due to lack of N-terminus interaction, highest protein-ligand interactions in CB_2_ macrostate *I^TM^*^17^ are observed in TM1 (Q32^1.31^) and TM7 (K279^7.33^, F283^7.37^, M286^7.40^). The anandamide also forms interactions with transmembrane lipids (phosphatidylcholine (PC), sphingolipids (SM), and cholesterol (CHL)) similar to the CB_1_ (Figures 4B).

Anandamide moves to the bound pose *B^Pro^_CB1_* from macrostate *I^TM^*^17^ (Figure 4C). In the macrostate, the major interactions with anandamide were observed with aromatic residues in TM2 (F91^2.61^, F94^2.64^), TM7 (F281^7.35^) and ECL2 (F183^ECL2^) (Figure 4C). K-L divergence analysis shows that this binding from macrostate *I^TM^*^17^ to *B^Pro^_CB1_* leads to the higher residue fluctuation in the N-terminus, extracellular TM1, and TM2 as ligand binds from the region (Figure 4D). Comparison of inactive structure of CB_1_ and CB_2_ points out less space between the TM2 and TM7 for CB_2_ along this pathway. To characterize major conformational changes in anandamide binding, F91^2.61^ *χ*_1_ dihedral angle and TM2 (F91^2.61^) TM7 (S285^7.39^) distance were plotted against each other (Figure S11A). MSM weighted free energy landscape reveal that F91^2.61^ *χ*_1_ angle ensemble average does not change as the ligand moves from macrostate *I^TM^*^17^ to *B^Pro^_CB1_*. However, the TM2 and TM7 distance increases to facilitate the anandamide binding (Figures S11A and S11B). Comparison of similar distances for CB_1_shows that TM2 and TM7 remain far away to be affected by ligand binding (Figure S12). Angle rotation of F174^2.61^ is enough for anandamide to move inside the pocket (Figure S12). As the helical movement is needed for CB_2_, the kinetic transition from microstate *I^TM^*^17^ to *B^Pro^_CB1_* is much slower (300.2 ± 99.8*µ*s) compared to the equivalent transition in CB_1_ (macrostate *I*1*^TM17^_CB1_* to *I*2*^TM17^_CB1_*: 29.0 ± 3.8*µ*s) (Figures 3C and 4C).

These analyses highlight the differences in the binding mechanisms of anandamide to CB_1_ and CB_2_ despite of binding from same the lipid access channel between TM1 and TM7. In CB_1_, the binding pocket shape changes during ligand binding due to the repositioning of the N-terminus, indicating an induced fit mechanism. Conversely, in CB_2_, the shape of binding pocket remains largely unchanged, suggesting a conformation selection mechanism. This demonstrates that endocannabinoids can utilize different binding mechanisms even when interacting with receptors from the same subfamily.

### Alternative anandamide binding pathway for CB_2_ in between TM5 and TM6

Another membrane-embedded ligand-binding pathway was detected for CB_2_ between TM5 and TM6. VAMPnets analysis shows two macrostates along this pathway before it converges back to the bound state (Figure 4C). In the macrostate closer to the membrane bilayer, the ligand is stabilized between the surface of TM5 and TM6. Diffusion from the membrane bilayer to *I*1*^TM^*^56^ is also relatively fast (3.4 ± 0.4*µ*s). This macrostate also represents a metastable region as the ligand is stabilized by the residues in TM5 (L195^5.44^) and TM6 (W258^6.48^, M265^6.55^, L259^6.59^) surface (Figures 4A and 4C). In previous experimentally determined structures, long chain lipid molecules were found to be bound to TM5 and TM6 surface, supporting our observation^16,45^ (Figure S13). The ratio of the protein and lipid surface area calculations indicates stronger interactions with CB_2_ surface as more than 80% of the ensemble averaged surface area embedded to the protein (Figure S6). Lipid interactions in this macrostate are limited to the polar head group of the anandamide as the hydrophobic tail remains inside the receptor (Figure 4B). K-L divergence analysis compared to macrostate *I*1*^TM^*^56^ shows a higher divergence in TM5 and TM6 surface to accommodate the tail of anandamide (Figure S14). Further analysis reveal that the pocket volume between the TM5 and TM6 interface of the CB_2_ increases as the ligand diffuses from the membrane to this intersurface (Figure S15).

Anandamide moves inside the receptor to macrostate *I*2*^TM56^_CB2_* from macrostate *I*1*^TM56^_CB2_*. Lack of lipid interactions confirms the ligand position inside the protein in this macrostate (Figure 4B). However, the ligand pose is different than compared to macrostate *B^Pro^_CB1_*. The major protein-ligand interactions were observed between the TM3 (I110^3.29^, T114^3.33^), TM5 (W194^5.43^), TM6 (W258^6.48^) (Figures 4A and 4C). Due to the stable interactions, RMSD calculations show that the ligand stabilizes in one stable pose in this macrostate (Figures 4C and S16). K-L divergence analysis shows that the major fluctuation happens in the TM6 due to the transition between macrostate *I*1*^TM^*^56^ to *I*2*^TM56^_CB2_* (Figure 4D). Major dynamic change was observed in the angle distribution of F202^5.51^ rotational movement to create space between TM3, TM5, and TM6 for the ligand. Anandamide bound in this macrostate increases the population of F202^5.51^ in the outward facing direction (Figure S17). Interstate transition further in the pocket between macrostate *I*2*^TM56^_CB2_* and *B^Pro^_CB1_* was also observed. Although anandamide binding is possible from both the lipid access channels for CB_2_, kinetic comparison between the pathways shows that pathway between TM1 and TM7 is kinetically more favorable (Figure 4C).

MD simulations were able to capture anandamide TM5 and TM6 for CB_1_, however, no metastable minima or ligand binding pathway were observed from this lipid access channel (Figure 2A). We explained this difference between CB_1_ and CB_2_ by calculating the lipid accessible surface area for the membrane-facing residues in the TM5 and TM6 surfaces (Figures S18A). Higher accessible surface area would increase the contact probability between anandamide and receptor to stabilize the ligand on the surface. For CB_2_, area per residue is much higher compared to the CB_1_ (Figure S18B). Less per residue surface area for CB_1_ can be attributed to bulky F368^6.60^ residue in the protein surface instead of the A270^6.60^ in CB_2_ (Figure S18A). This decreasing surface area implies that the anandamide cannot reach the groove between TM5 and TM6, decreasing stable interactions between the ligand and CB_1_.

### Distinct binding pocket interactions contributes to anandamide selectivity

Difference in the binding mechanism leads to dissimilar binding poses and interactions of anandamide inside the pocket. Recently, cryo-EM structure of anandamide analog (AMG315) bound CB_1_ was determined in the active state with downstream G-protein.^46^ AMG315 is functionally a full agonist and is observed in the canonical agonist-bound pose in the cyroEM structure. Comparison of AMG315 bound structure with MD sampled structures of the bound poses shows that MD simulations are able to capture the experimentally determined ligand pose for both CB_1_ and CB_2_ (Figures S19A and S19B). However, distance RMSD calculations of anandamide in bound poses show that the canonical agonist bound conformation is not the most stable conformation for anandamide in the pocket (Figures S19C and S19D). We performed relative binding free energy (RBFE) calculation with respect to anandamide analogs with known experimental binding affinity to computationally validate these obtained distinct poses. One major drawback of this method of validation is that RBFE calculation depends on the reference ligand’s pose. However, anandamide shows significant flexibility inside the pocket for both receptors. There is high enough probability that ligand can obtain other poses in the receptor, which can affect the calculated ΔΔG. Hence, instead of the quantitative agreement, we focus on the qualitative trend of ΔΔG with respect to most stable pose of anandamide.

The analogs that we are studying are the ones where scaffold changes have been focused on the polar group of the anandamide, as polar groups form the most stable interactions in the pocket (Figure S20A). The selected analogs were classified into three categories to better understand the effect of scaffold change: (1) addition of alkyl groups in C1’ and C2’. It has been shown that adding an alkyl group in this position makes the ligand more resistant to hydrolysis by degrading enzymes, increasing its apparent binding affinity.^47^ (2) removal of terminal hydroxyl group. Previous studies have shown that the hydroxyl group is not essential for binding.^27^ (3) substitution of hydroxyl group with fluorine group. These analogs increase the CB_1_ and CB_2_ affinity.^27^ Estimated ΔΔG reveals that binding affinity increases for CB_1_ and CB_2_ for all three categories of the analogs and magnitudes of the ΔΔG also remain within the realistic limits (Figure S20B). Considering the assumption of RBFE calculations, these calculations validate the binding pose obtained from the simulations.

These distinct binding poses and differences in the receptors’ conformation due to ligand binding lead to difference of ligand interaction with receptors. To observe these differences, we compared the interactions of anandamide with equivalent loops and helices surrounding orthosteric pocket of receptors. Interaction energies were calculated using LIE method for 1000 representative structures from bound state of each receptor, which were selected based on their MSM probabilities. Ligand interaction energy calculations in these structures show major differences in N-terminus region of the two receptors (Figure 5A). In spite of extracellular motion, stronger interactions with N-terminus hydrophobic residues were observed for CB_1_ compared to CB_2_. Furthermore, overall comparison of enthalpy contribution due to protein and ligand interactions indicate higher enthalpy contribution for CB_1_. This makes the anandamide binding more enthalpically favorable to CB_1_.

**Figure 5:**
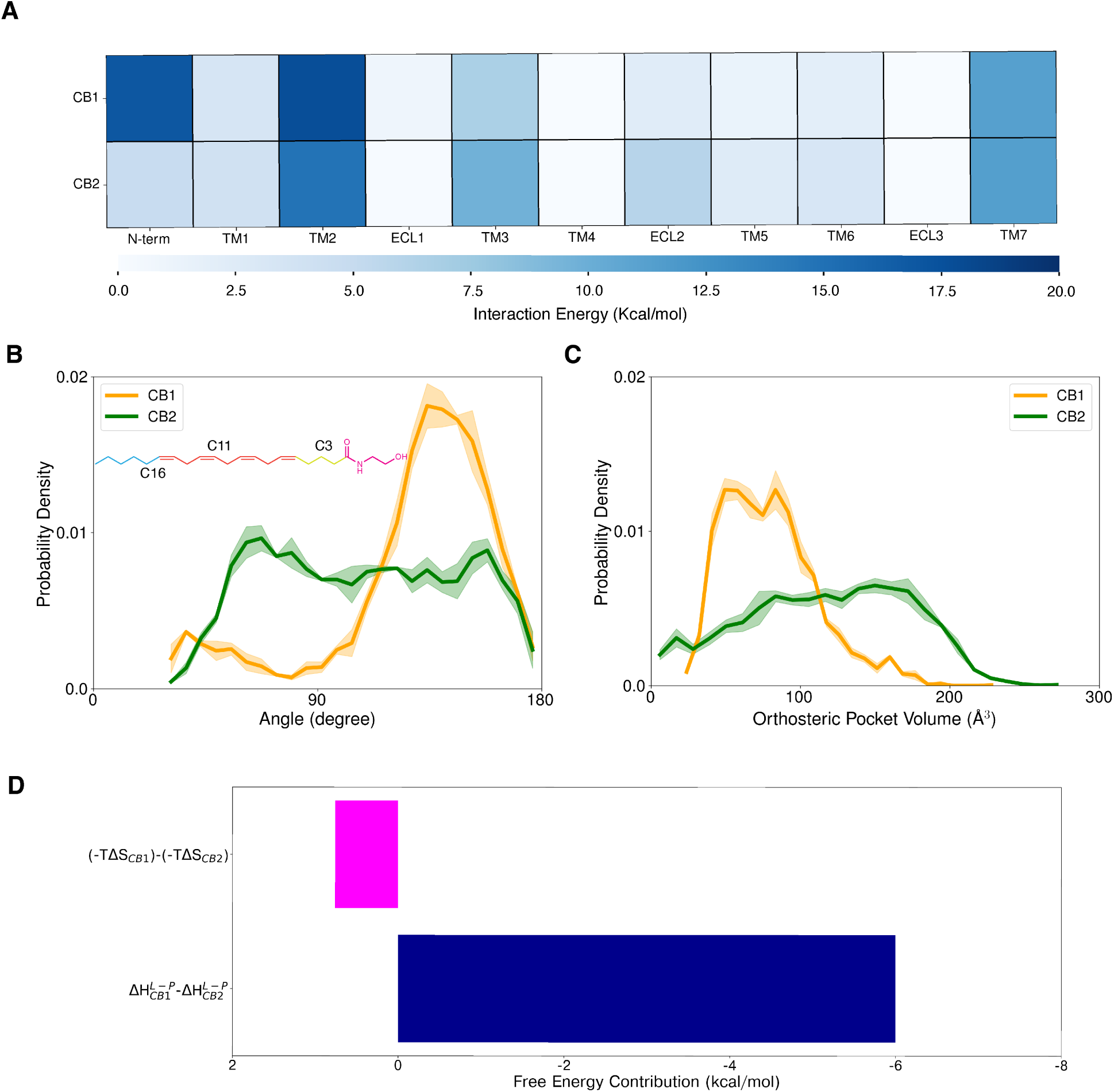
(A) Anandamide interaction energies of different structural elements of cannabinoid receptors in the bound states. (B) Probability density plot of three *sp*^3^ carbons of anandamide representing hydrophobic tail (C16), polyene linker (C11), and propyl linker (C3) for *B^Pro^* (color: orange) and *B^Pro^* (color: green). (C) Probability density plot of binding volume calculation of orthosteric binding pocket for *B^Pro^* (color: orange) and *B^Pro^* (color: green). Error in angle and pocket volume distributions were calculated from on 3 bootstrap samples where each sample contains 1000 frames obtained from macrostate based on MSM weighted probability. (D) Bar plot represents the differences of enthalpy (color: blue) and entropy (color: magenta) contribution between the *B^Pro^* and *B^Pro^*.

### Explanation of CB_1_ selectivity of anandamide

Although there are conflicting reports on the binding affinities of anandamide, the general consensus is that anandamide is more selective towards CB_1_ compared to CB_2_.^18^ We showed that the ligand binding is enthalpically favorable to CB_1_. However, significant ligand dynamics inside the pocket due to large pocket volume of receptors and plethora of sp^3^ hybridized carbon atoms may lead to a significant entropy effect in the binding free energy. (Figures S19C and S19D).

To compare the anandamide dynamics inside CB_1_ and CB_2_ pocket, we calculated angle between carbon atoms at three different parts of ligand: tail (C16), polyene linker (C11), and propyl linker (C3) group (Figure 5B). This analysis shows that anandamide is stabilized in one conformation for majority of the population in *B^Pro^_CB1_*. In contrast, the ligand has much higher divergence in angle distribution for CB_2_ with multiple equally probable angular conformations (Figure 5B). This higher conformational divergence in macrostate *B^Pro^_CB1_* can be explained by the binding pocket volume calculations (Figure 5C). These calculations show that anandamide has larger accessible volume to move in CB_2_ compared to the CB_1_, which matches with our previous observations.^38^ Therefore, in CB_1_, degrees of freedom for the movement of sp^3^ hybridized carbons are restricted compared to CB_2_. The higher fluctuation in ligand due to larger volume in CB_2_ leads to higher entropic contribution in free energy (Figure 5D). Estimation of Gibbs entropy (Δ*S* = − ^L,^*_i_ k_B_p_i_ln*(*p_i_*)) show higher entropic contribution for anandamide binding in CB_2_, which contradicts with the higher enthalpic effects in CB_1_. Comparison of these contrasting effects of enthalpy and entropy between two receptors, enthalpy contribution to free energy dominates in the ligand binding process (Figure 5D). Hence, anandamide binds favorably to CB_1_ than CB_2_.

## Conclusions

Endocannabinoids are lipid-like long-chain molecules that bind to cannabinoid receptors to transduce intracellular signaling. One of the endocannabinoids, anandamide, has garnered significant attention due to its partial agonistic nature and CB_1_ selectivity. Designing analogs of anandamide as CB_1_ selective drugs necessitates a deep understanding of its binding mechanisms. In this study, we developed two hypotheses to elucidate the CB_1_ selectivity of anandamide.

First, we proposed that structural differences along the ligand-binding pathway lead to distinct bound poses and the resulting enthalpy contributions to the ligand-binding free energy. Second, we suggested that the distinct pocket volumes of CB_1_ and CB_2_ lead to differences in ligand dynamics, influencing the entropy contributions to free energy. To test these hypotheses, we conducted unbiased extensive MD simulations of anandamide binding to CB_1_ and CB_2_.

Our MD simulations reveal mechanistic differences in anandamide binding between CB_1_ and CB_2_ via the pathway between TM1 and TM7. Binding to CB_1_ is characterized by an induced fit mechanism, where the ligand displaces the N-terminus from the orthosteric pocket. In contrast, CB_2_ binding suggests a conformation selection mechanism, with no major changes observed. For CB_2_, a separate non-canonical binding pathway was observed in between TM5 and TM6. Although this pathway finally converges to the same orthosteric binding pocket, it is kinetically unfavorable compared to the canonical pathway between TM1 and TM7. These distinct binding mechanisms result in different ligand binding poses and interactions with the receptors. Major differences in ligand interactions are observed between the N-terminus of the receptors. Higher interaction with N-terminus for CB_1_ makes the ligand enthalpically more stable in CB_1_ binding pocket, which supports our first hypothesis. Additionally, we observed that the larger pocket volume of CB_2_ allows greater ligand dynamics, leading to a more favorable entropy contribution to free energy for CB_2_. Comparison of this relatively contradicting enthalpy and entropy effect on binding free energy reveal that the dominant enthalpy effect causes the ligand to be more selective towards CB_1_. In summary, our work elucidates the orthosteric selectivity of anandamide for cannabinoid receptors using extensive molecular simulations and post-processing Markovian techniques. These insights provide crucial information for designing selective drugs.

## Methods

### System Preparation

Inactive structures of CB_1_ (PDB ID: 5TGZ^15^) and CB_2_ (PDB ID: 5ZTY^16^) were selected to perform anandamide binding simulations. Non-protein residues (ligands and water molecules) and fused proteins (Flavodoxin and T4-lysozyme) were removed for the structure file. Thermostabilized mutations were modified back to the original residues. Missing residues in ICL3 were modeled using Rosetta loop modeling for CB_1_ and CB_2_.^48,49^ For CB_2_, missing 21 residues in the N-terminus were also modeled using Robetta webserver.^50^ Ninety-eight residues were missing from the CB_1_ N-terminus, and no template is available to model from homologous protein.^51^ Furthermore, it is challenging to model 98 residues accurately with template-free modeling because of the combinatorial expansions of conformational space.

However, the membrane-proximal regions of the N-terminus were shown to be important for CB_1_ signaling by allosterically modulating ligand affinity.^52^ Hence, the closest ten residues (89 to 98) were modeled using Rosetta with a disulfide bond constraint between C98^N−term^ and C107^N−term^ residues.^52,53^ The remodeled structure with the least energy was further refined in each step using kinematic closure protocol. ^54^

Proteins with the modeled loops were embedded in the membrane bilayer and salt solution (extracellular and intracellular region) using CHARMM-GUI.^55^ Asymmetric complex membrane compositions were selected for CB_1_ and CB_2_. Average brain membrane composition was used for CB_1_ as it is majorly expressed in the central nervous system. Average mammalian cell composition was used for CB_2_ as it is majorly expressed in immune cells. The membrane compositions were obtained from Inǵolfsson et al. and proportionally downsized according to our system.^56^ The final membrane composition is shown in Tables S1 and S2. 150 mM NaCl salt solution with TIP3P water model is used in the extracellular and intracellular regions.^57^ CHARMM36m forcefield was used to parameterize the system.^58^

Additional steps were implemented to include anandamide in MD systems. Anandamide was parameterized with CGenFF using the “Ligand Reader & Modeler” of CHARMMGUI.^59^ PACKMOL v18.169 software package randomly placed the anandamide in the extracellular solution.^60^ Ligand added systems were rebuilt using the psfgen v2.0 module of VMD.^61^ ParmEd v3.2.0 software package was used to perform hydrogen mass repartitioning on the systems and convert the systems into AMBER-recognized format.^62–64^

### Simulation Details

Minimization, equilibration, and initial round of production runs were performed in AMBER v18 simulation package.^65^ Later rounds of production runs were run on openMM v7.7 using the distributed computing platform folding@home.^66,67^ Systems were minimized for a total of 14000 steps with an initial 5000 steps using the steepest descent algorithm and the remaining 9000 steps with conjugate gradient algorithm.^68^ Temperature was increased to 300K in NVT ensemble using Langevin dynamics as thermostat.^69^ Collision frequency for Langevin dynamics was used as two ps^−1^. NVT ensemble simulations were performed for one ns. In the next step, the pressure was set to 1 bar in the NPT ensemble using a Monte Carlo barostat with anisotropic pressure control.^70^ This step was performed for two ns. In NVT and the initial stage of NPT ensemble simulation, the protein backbone atoms were restrained using a harmonic force with spring constant 10 kcal/mol-Å^2^. These steps were run using the AMBER v18 module sander. The restraints were removed in the final equilibration step, and systems were equilibrated for 50 ns in the NPT ensemble before the production run. Production runs were also continued in the NPT ensemble. During the simulation, a cut-off distance of 12 Å was used for non-bonded force calculations. Particle Mesh Ewald method was used for long-range electrostatic calculations.^71^ SHAKE algorithm was used to constraint bond lengths for bonds with hydrogen atoms.^72^ As HMR was used to perform simulations, four fs simulation step was chosen.^63,64^ Periodic boundary conditions were maintained throughout the simulation. All the simulation parameters were kept the same during AMBER v18 and openMM v7.7 production runs.

### MD sampling

As GPCR orthosteric pockets are buried inside the transmembrane domain, ligand binding often involves stable intermediate states. ^23,73–76^ Therefore, long timescale MD simulations are not efficient for capturing ligand binding as it requires significant time to sample the entire process. Here, we employed adaptive sampling,^77,78,78–86^ a exploitative and explorative iterative approach for sampling the ligand binding by running parallel short simulations. Adaptive sampling has been successfully employed to investigate the dynamics of mmebrane proteins including GPCRs.^23,38,75,87–92^ Initial rounds of sampling were exploitative, where we ran simulations from the frames with ligand positions closest to the binding pocket. The distance of the ligand center of mass from the protein binding pocket center of mass was estimated along the y and z projection, and the closest 15 to 20 frames were selected to run the next round of sampling. Residues to calculate the center of mass of protein binding pocket for CB_1_ and CB_2_ are shown in Figure S21. Each trajectory was run for 80 ns. 28 and 25 rounds of iterative simulations were run for CB_1_ and CB_2_ to observe the complete binding of the ligands.

As MSM building requires sampling of reversible local transitions during the binding procedure, further sampling was done from the entire simulation space using folding@home distributive computing.^66^ The entire simulation space was clustered into 1000 clusters based on x, y, and z projection of the center of mass (COM) distance from the ligand. One frame was picked randomly from each cluster, and four independent 100 ns atomistic unbiased simulations were run using folding@home distributive computing.

Finally, explorative sampling was performed to sample the ligand diffusion in the membrane bilayer. Simulations were started from the frames where ligand exists in the membrane. For each round of simulations, these frames are clustered into 20 clusters based on x, y, and z projection of the center of mass (COM) distance from the ligand, and one frame is selected randomly from each cluster. From the selected frame, four independent simulations were started using folding@home. Simulations were continued until the membrane adjacent area of the protein was appropriately sampled.

### Featurization of Protein Conformational Ensemble

To build Markov state model (Discussed in the next subsection), each conformation of the biomolecular simulations needs to be featurized to capture the important dynamics. Here, ligand binding and resulting protein conformational changes are the process of interest. To featurize the ligand binding dynamics, the distance from the hydrophobic tail’s terminal carbon (C20) and anandamide’s carboxyl oxygen to every binding pocket residue (C*α*) was calculated. For capturing protein movements, we considered the pairwise residue distances of residues that undergo significant conformational change during activation as measured by the residue-residue contact scores (RRCS).^93^ The entire process of residue pair selection using RRCS is discussed in Dutta and Shukla.^38^

### Markov State Models

As short trajectories were run to observe ligand binding, each trajectory can only sample the local transitions. Markov state model (MSM) is a technique that connects the local transitions observed in short trajectories to provide global information about the dynamics such as free energy barriers and kinetics of transitions.^94–98^

In MSM, the MD simulations are discretized into microstates. It is assumed these states would follow Markovian dynamics where the jump between states would only depend on the current state, not the past state.^99^ A concept of lag time was introduced to maintain this Markovian property between interstate transitions. Lag time is the minimum time when the interstate transitions become memoryless. Lag time (*τ*) for CB_1_ and CB_2_ ligand binding systems were considered 35 ns (Figures S22A and S22B).

According to the assumption of Markovian property, all conformations in a macrostate have to be kinetically similar. Therefore, before discretizing the simulation spaces, featurized MD frames were converted into time-independent components (tiCs) using the variational approach of MSM.^100,101^ tiCs are the linear combinations of slow features. Here, tICs were built with a lag time of 25 ns. tiCs were further clustered, and transitional probabilities (*p_ij_*) between the clusters were calculated using maximum likelihood estimation.

To build an optimized model, hyperparameters such as the number of tIC components and microstates were optimized by maximizing the VAMP-2 score.^102^ VAMP-2 scores are a measure to represent the quality of the slowest processes captured by Markov state models (Figures S22C and S22D). In this case, the VAMP-2 score was calculated using a 10-fold cross-validation of the MD data. The entire MSM building process was implemented using pyEMMA 2.5.6 python package.^103^ Optimized MSM redistributes the population of Markovian states, removing the bias created by running short simulations (Figures S23A and S23B).

### Trajectory Analysis and Visualization

Feature calculations (e.g., distances, angles) for MSM building and other analyses were performed using MDTraj 1.9.8 python package.^104^ Trajectory visualization was performed VMD v1.9.3.^61^ Frame selection and other modulations in trajectories were performed using CPPTraj v18.01. All analysis codes were written using written in python programming language. Pocket volume calculations were performed using POVME3.0.^105^ Getcontacts software was used for contact probability calculations.^106^ Linear interaction energy (LIE) analysis was performed using CPPTraj implementation to calculate the interaction energies between the ligand and protein residues.^107,108^

### Metastable State Estimation Using VAMPnets

VAMPnets is a deep learning based technique to capture the slow processes from MD ensemble.^41^ Using neural network architecture, VAMPnets tries to find the non-linear transformation of features that can represent slow processes. VAMPnets is trained by maximizing the VAMP-2 score. In this work, VAMPnets were used to discretize the space into macrostates. Macrostate represents a collection of Markovian states that encompasses kinetically close regions of the MD ensemble.^38^ To achieve the discretization, we used the softmax activation function at the last layer of the neural network. Softmax function returns a discrete probability distribution, where each value represents a probability of an MD frame belonging to a particular macrostate.

We used features that have been used to build MSM to train VAMPnets. A forward neural network with eight hidden layers was used for training, where number of nodes per layer is 30, 100, 100, 100, 100, 100, 100, 30, respectively. ELU activation was used on the input of every hidden layer. The network was trained for 100 epochs on 80% of total trajectories.

The number of the macrostates for every system was selected based on the least errors in the implied time scale and macrostate population^109^ (Figures S2, S3, S4, and S5). Implied time scale is obtained from the VAMP transformation of the last layer. The VAMPnets analysis was performed with Deeptime 0.4.3 python package.^110^

### Transition Path Theory

Transition Path Theory (TPT) is a probabilistic approach to calculate the kinetics from the Markov state model.^111,112^ This approach introduces a variable named committer (*q*^+^). In a probabilistic transition between an initial and final macrostate, committer (*q*^+^) is the probability of an intermediate Markovian state (*i*) reaching the final state before the initial state. Mathematically, the committer can be calculated from the system of linear equations as shown in Equation 1. From the committer, the mean free passage time between the two macrostates is calculated using Equation 2, where *π_i_* is the stationary density of *i*th Markovian state and *τ* is the lagtime of MSM. TPT calculations were performed using PyEMMA v2.5.6.^103^ Error estimation in TPT calculations was performed using the bootstrapping approach, where MSM was built using randomly selected 80% of all trajectories.^113^

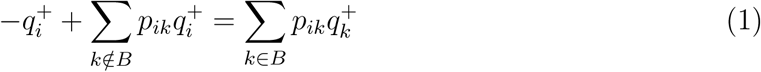

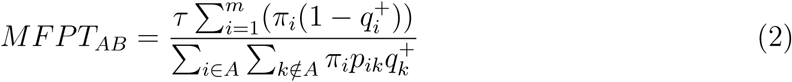

### Kullback-Leibler Divergence

Kullback-Leibler divergence (K-L divergence) is an asymmetric statistical distance measurement between two probability distributions. In this context, K-L divergence was performed between the distance distributions of difference macrostates to compare protein conformational changes observed during the ligand binding process.^23,38,114^ For this analysis, a thousand representative conformations from each macrostate were selected based on the probability of the MSM states belonging to the macrostate. All possible pairwise closest-heavy atom distances (except for the two next neighbors per residue) were calculated for these conformations of each macrostate. K-L divergence was estimated between the inverse of each pair of these distances. Per residue pair, we averaged two measurements where each macrostate is considered a reference to make it symmetric. To estimate per-residue contribution, we averaged K-L divergences for all residue pairs containing that certain residue.

### Relative Free Energy Calculation

We selected 11 synthetic analogs of anandamide to perform relative binding free energy calculations on the stable bound pose of anandamide observed in the simulation. These calculations were performed on the GROMACS v2020.7 simulations package with a single topology alchemical free energy approach.^115^ System building was performed with the following steps.

- As anandamide and its analogs are flexible, a constrained optimization module of the RDKit was used to generate the coordinates of analogs from isomeric smiles stings similar to the bound pose of anandamide in CB_1_ and CB_2_. The isomeric smiles were obtained from the PubChem database.^116^
- OpenBabel v2.4.1 software was used to convert the obtained mol file from RDKit to mol2 format and to add hydrogen.^117^ The mol2 file was parameterized using the CHARMM-GUI ligand generator module. The ParmEd package was used to create the top and gro files.
- alchemical-setup module was used to create single topology top and gro files of the analogs by mapping the similar atoms against the anandamide.
- The new ligand system was solvated using a gmx module of GROMACS, and a 150 mM concentration of NaCl was maintained.
- To create the complex system, the anandamide-bound receptor system was converted into the top and gro format using ParmEd software. Further, anandamide topology parameters and coordinates were replaced with the ligand system.

To perform Alchemical free energy calculations, the system is divided along a hyperparameter *λ*, where *λ* equal to 0 represents the reference ligand and *λ* value 1 represents query ligand.^118^ Here, twenty-one *λ* values were considered for both ligand and complex systems. In Intermediate *λ* values, force-field parameters were obtained by the linear combination of both ligand parameters.

Ligand systems and complex systems were minimized and equilibrated using NVT and NPT ensemble. The steepest descent algorithm was used for energy minimization for 15000 steps. Langevin dynamics was used as a thermostat to maintain the temperature at 300K.^69^ Parrinello-Rahman barostat was used to control the pressure at 1 atm.^119^ Cutoff distances for electrostatics and other non-bonded interactions were 12 Å. The Particle Mesh Ewald method was used for long-range electrostatic calculations. To take into account the particle disappearance, the soft-core version of the potential was used to with hyperparameter *σ*, *α* and soft-core *λ* power set to 0.3, 0.5, and 1 respectively.^120^ Each NVT and NPT step was run for 100 ps. For a complex system, the backbone atoms of the protein are restrained with 1000 kJ/mol. Production runs for each *λ* value of each complex and ligand system were performed for five ns. Free energy calculations from these simulations were performed with a thermodynamics integration approach using the Python module alchemical-analysis.^121^

## Acknowledgements

D.S. acknowledges the support of NSF Early CAREER Award (MCB-1845606) and NIGMS MIRA award R35GM-142745 for this project. The authors thank the volunteers of folding@home for donating computing time. S.D. thanks Austin Weigle for the discussion during system setup.

## Author contributions

D.S. obtained funding for the project. D.S. supervised the research. D.S. and S.D. designed the research. S.D. and L.Z built the MD systems and performed simulations. D.S. participated in discussing results. S.D. analyzed data, and wrote the article with inputs from D.S. All authors reviewed the manuscript.

## Code availability

Python scripts to reproduce the results are publicly available at GitHub. https://github.com/ShuklaGroup/Dutta_Shukla_Cannabinoid_2023b.git

## Data availability

Simulation topology, trajectory files, MSM objects and other necessary files to reproduce the analyses shown in the manuscript can be obtained from https://uofi.box.com/s/q9w1f99mybxo6hrgo8sfr3sbm3l8xrmu.

