## Supplementary Information for "Dynamic Mechanism for Subtype Selectivity of Endocannabinoids"

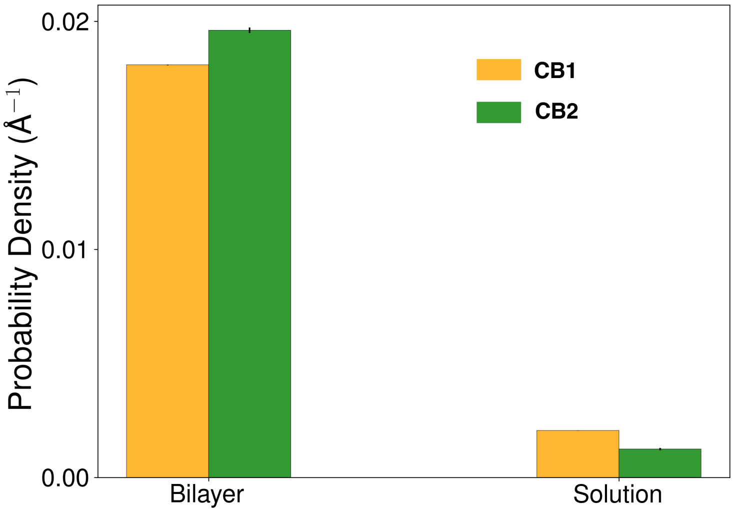


Figure S1: Bar plots show the probability densities of anandamide in membrane bilayer (excluding protein) and solution for $CB_1$ (color:Orange) and $CB_2$ (color:green). Error bars were calculated from three bootstrap samples, where MSM was built on sample with randomly selected 80% of total trajectories.

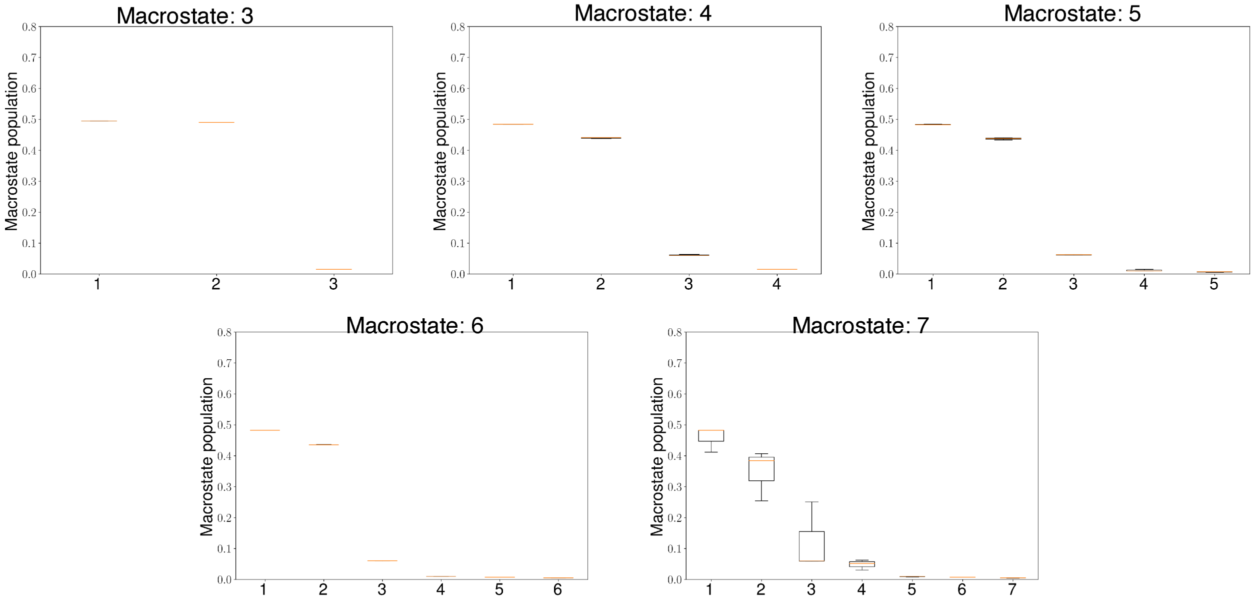


Figure S2: The box plots show the population of the $CB_1$ macrostates obtained by training VAMPnets. VAMPnets were trained based on different numbers of nodes in the output layer, which correspond to different numbers of macrostates for anandamide binding simulations of $CB_1$. VAMPnets with particular macrostates were trained 3 times 80% of total trajectories selected randomly. Population of each macrostate is equal to the sum of stationary density of all the microstates belonging to that macrostate.

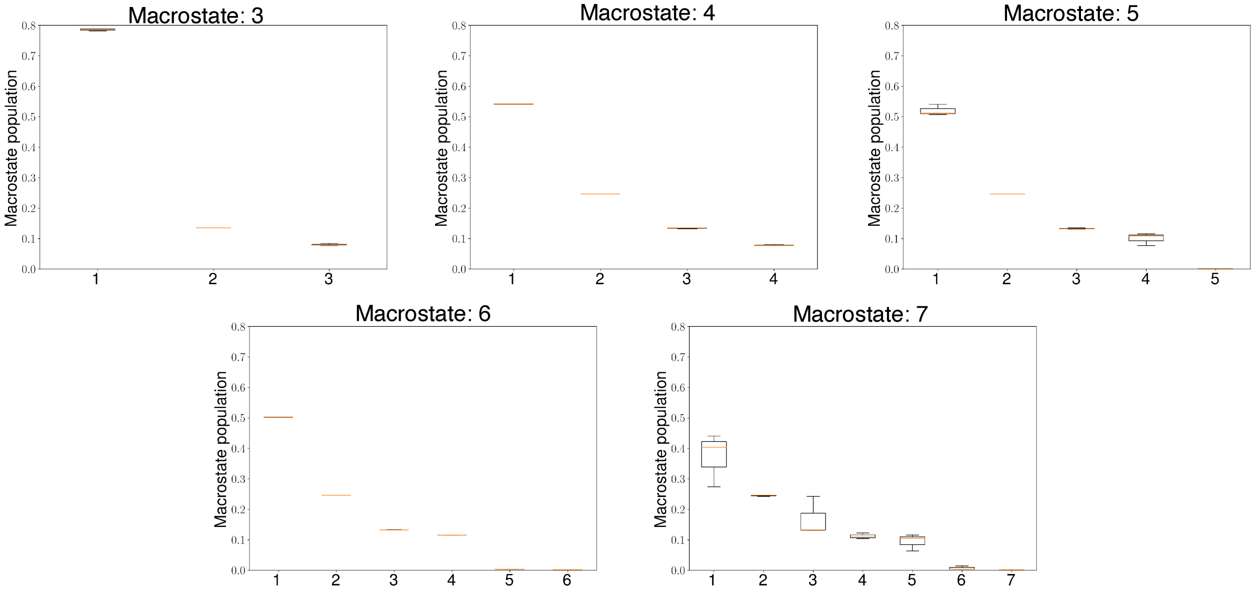


Figure S3: The box plots show the population of the $CB_2$ macrostates obtained by training VAMPnets. VAMPnets were trained based on different numbers of nodes in the output layer, which correspond to different numbers of macrostates. VAMPnets with particular macrostates were trained 3 times 80% of total trajectories selected randomly. Population of each macrostate is equal to the sum of stationary density of all the microstates belonging to that macrostate.

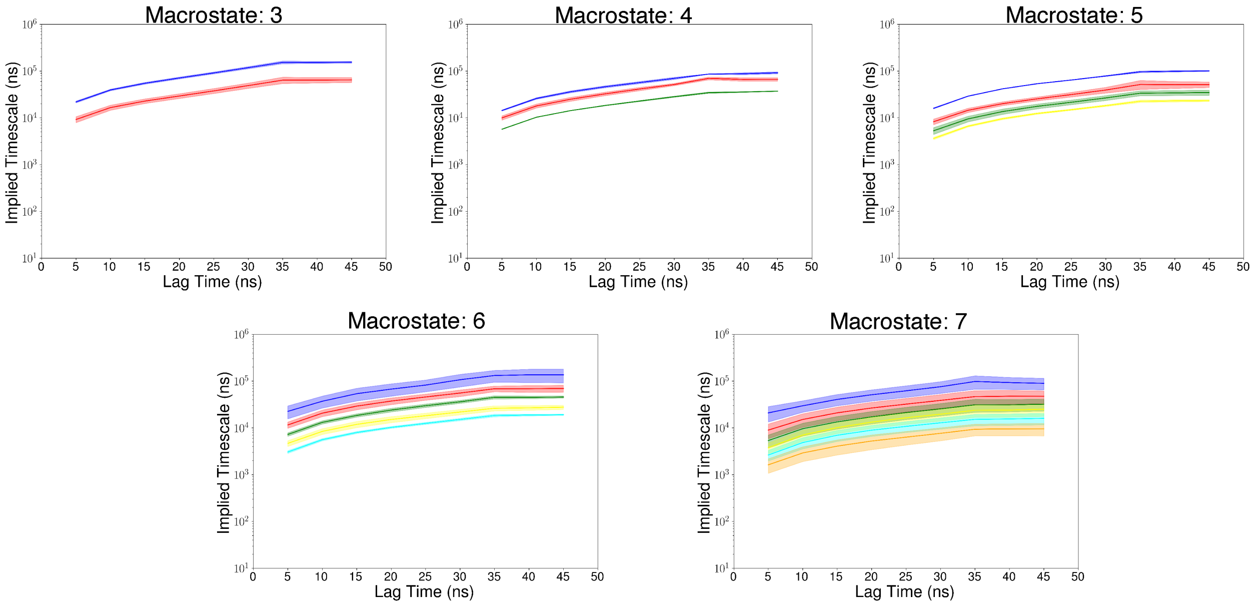


Figure S4: Implied timescale for inter-macrostate jumps are plotted against lag time for systems with different number of macrostates for anandamide binding simulations of $CB_2$. Implied timescales were calculated from the VAMP transformations of the last layer of the VAMPnets. VAMPnets with particular macrostates were trained 3 times 80% of total trajectories selected randomly.

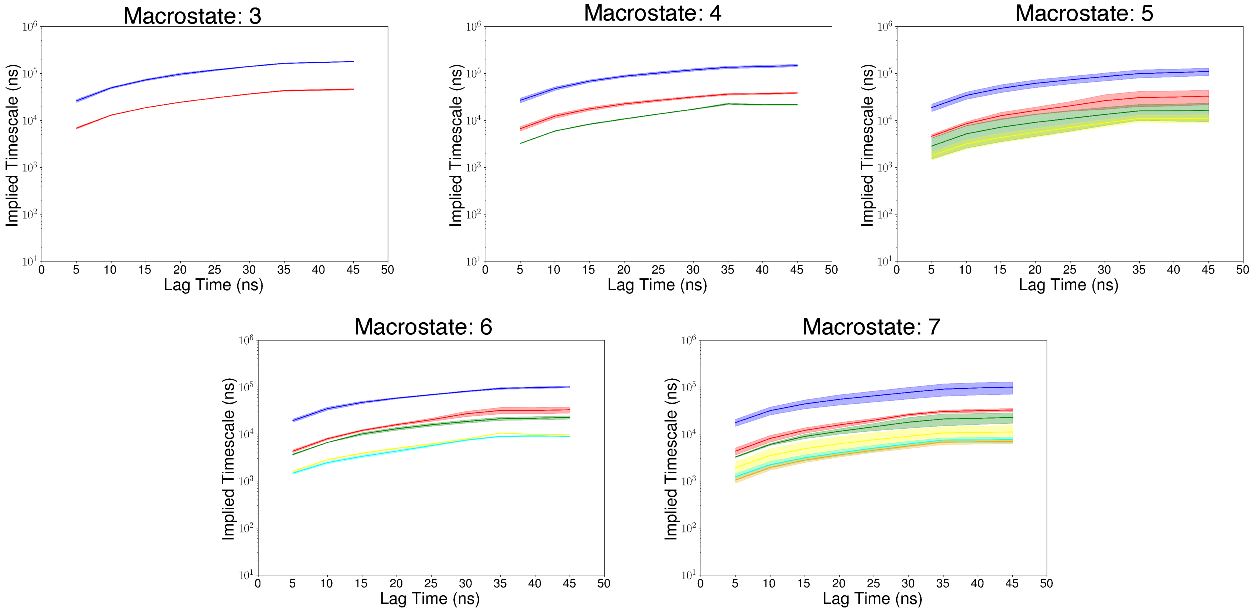


Figure S5: Implied timescale for inter-macrostate jumps are plotted against lag time for systems with different number of macrostates for anandamide binding simulations of $CB_1$. Implied timescales were calculated from the VAMP transformations of the last layer of the VAMPnets. VAMPnets with particular macrostates were trained 3 times 80% of total trajectories selected randomly.

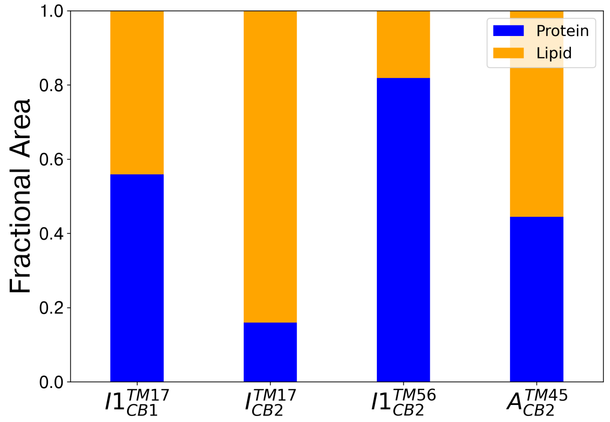


Figure S6: Ratio between protein (color: blue) and lipid (color: orange) exposed surface area of anandamide in different macrostates ($I1_{CB1}^{TM17}$, $I_{CB2}^{TM17}$, $I1_{CB2}^{TM56}$, $A_{CB2}^{TM45}$) where ligand stay in protein-membrane interface. Ratio was obtained by averaging surface area calculations from 3 bootstrap samples of each macrostate which consists of 1000 frames obtained based on MSM weighted probability.

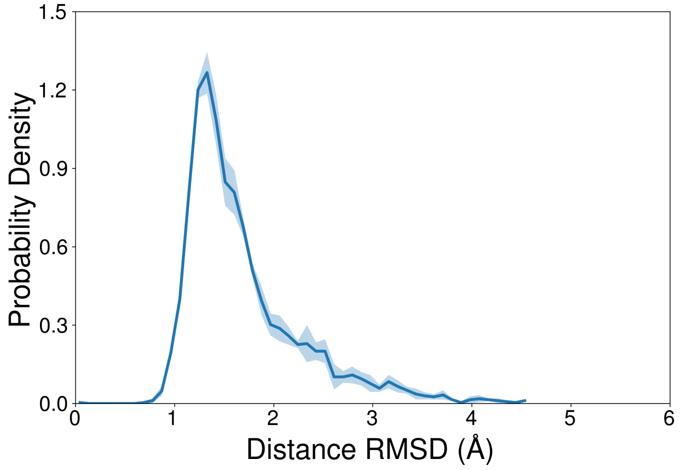


Figure S7: Probability density plot of distance RMSD for $I1_{CB1}^{TM17}$ macrostate. Error in distance RMSD calculations were calculated from on 3 bootstrap samples where each sample contains 1000 frames obtained from macrostate $I1_{CB1}^{TM17}$ based on MSM weighted probability.

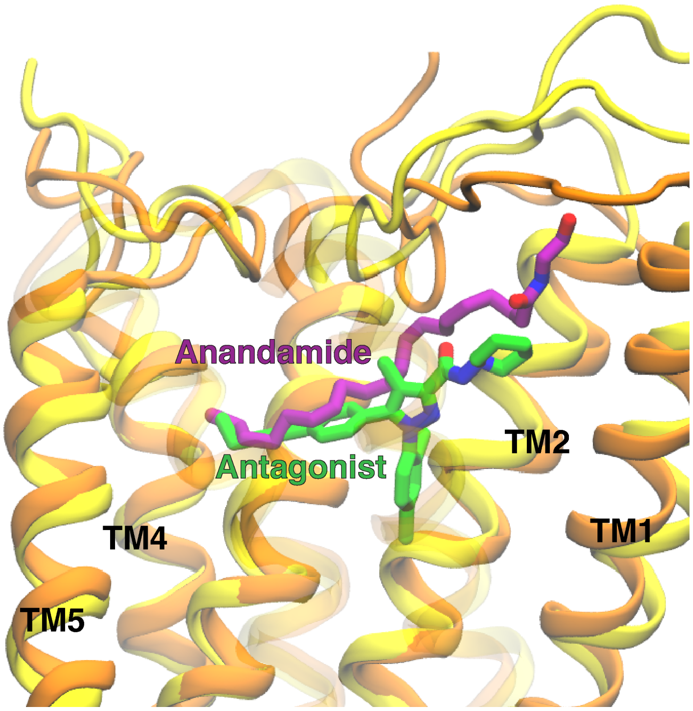


Figure S8: Superposition of antagonist (AM6538) bound inactive structure (PDB: 5TGZ,[1] color: orange) and representative structure from $I2_{CB1}^{TM17}$ macrostate (color: yellow). Ligands (Anandamide: purple; AM6538: green) and proteins are shown as sticks and cartoon representations. Transparent representation of TM6 and TM7 are used to show the ligand binding position.

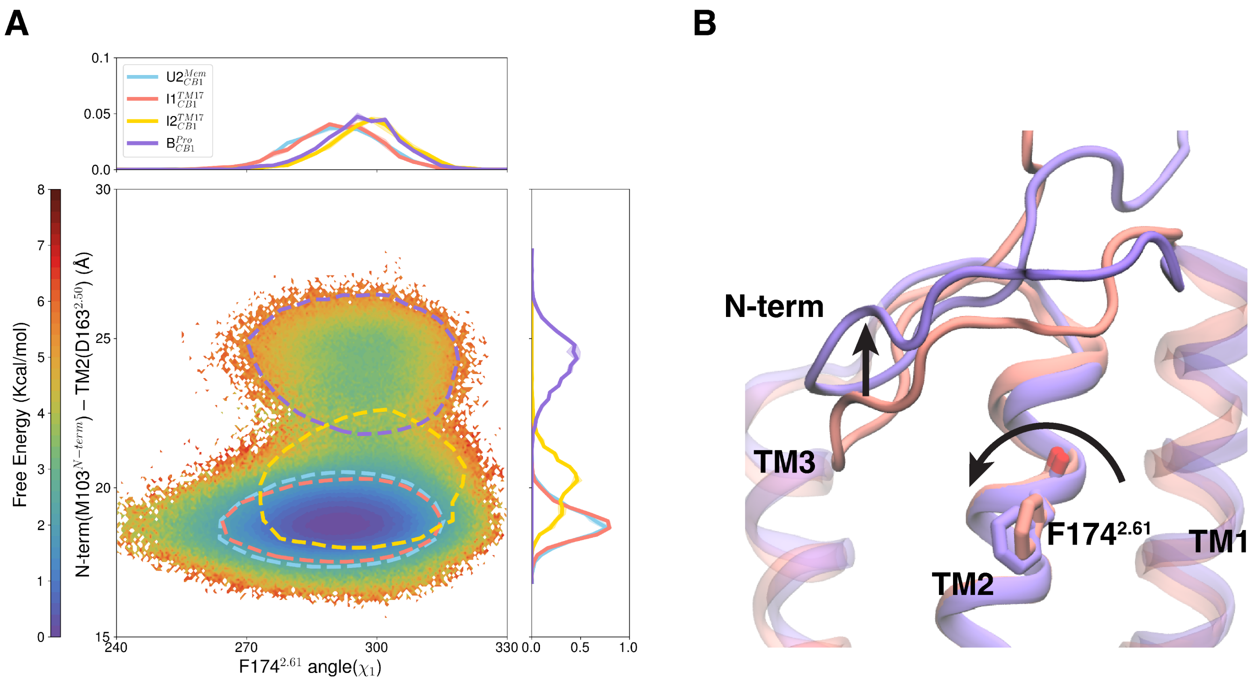


Figure S9: (A) For $CB_1$, MSM weighted free energy landscape free energy landscape is plotted between N-terminus - TM2 distance and F174$^{2.61}$ $\chi_1$ angle. Projections of important macrostate positions are shown as contour line on the landscape. Each feature of the free energy landscape is also plotted as probability density plot for different macrostates. (B) Representative structures from macrostate $I1^{TM17}_{CB1}$ (color: salmon) and $B^{Pro}_{CB1}$ (color: mediumpurple) of $CB_1$ are superposed. Important conformation changes from macrostate $I1^{TM17}_{CB1}$ to $B^{Pro}_{CB1}$ are shown with arrow direction.

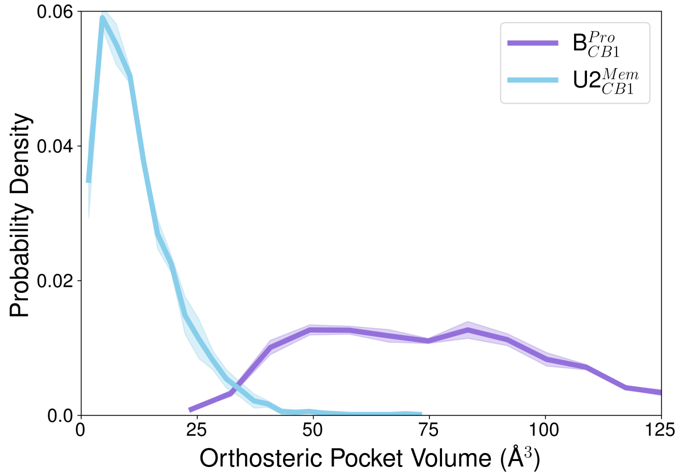


Figure S10: Probability density plot of binding volume calculation of orthosteric binding pocket for $B^{Pro}_{CB1}$ (color: mediumpurple) and $U2^{Mem}_{CB1}$ (color: skyblue). Error in angle and pocket volume distributions were calculated from on 3 bootstrap samples where each sample contains 1000 frames obtained from macrostate based on MSM weighted probability.

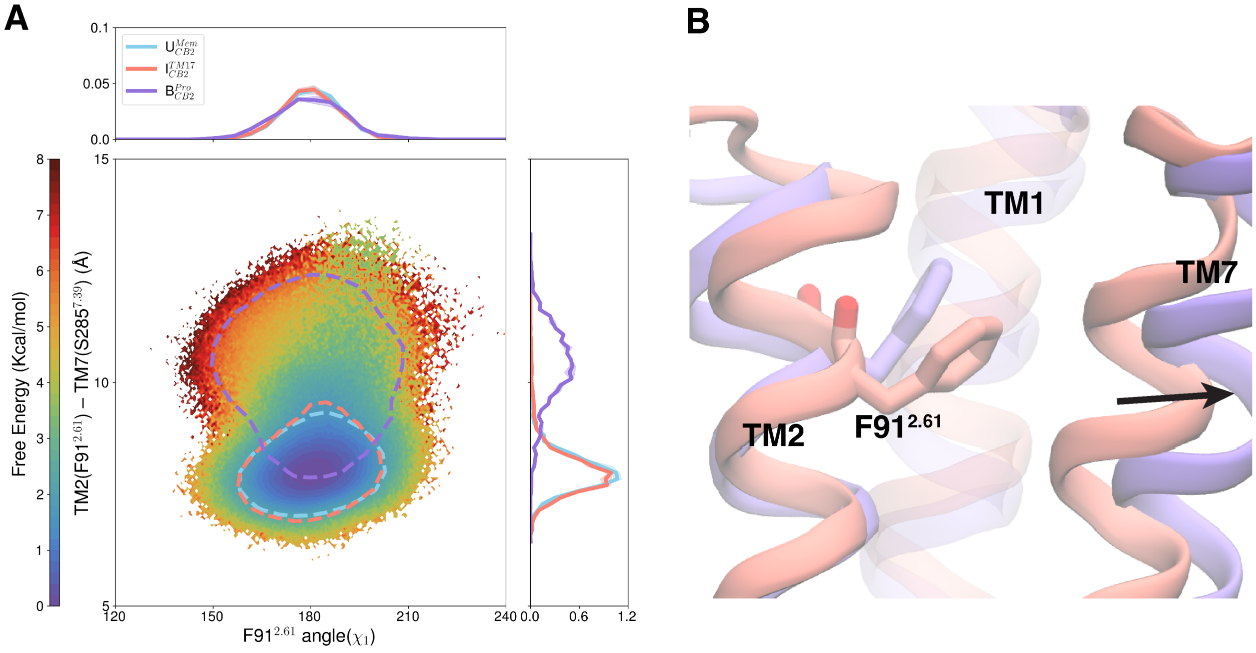


Figure S11: (A) For $CB_2$, MSM weighted free energy landscape free energy landscape is plotted between TM2 - TM7 distance and F91$^{2.61}$ $\chi_1$ angle. Projections of important macrostate positions are shown as contour line on the landscape. Each feature of the free energy landscape is also plotted as probability density plot for different macrostates. (C) Representative structures of macrostate $I^{TM17}_{CB2}$ (color: salmon) and $B^{Pro}_{CB2}$ (color: mediumpurple) of $CB_2$ are superposed. Important conformation changes are shown with arrow direction.

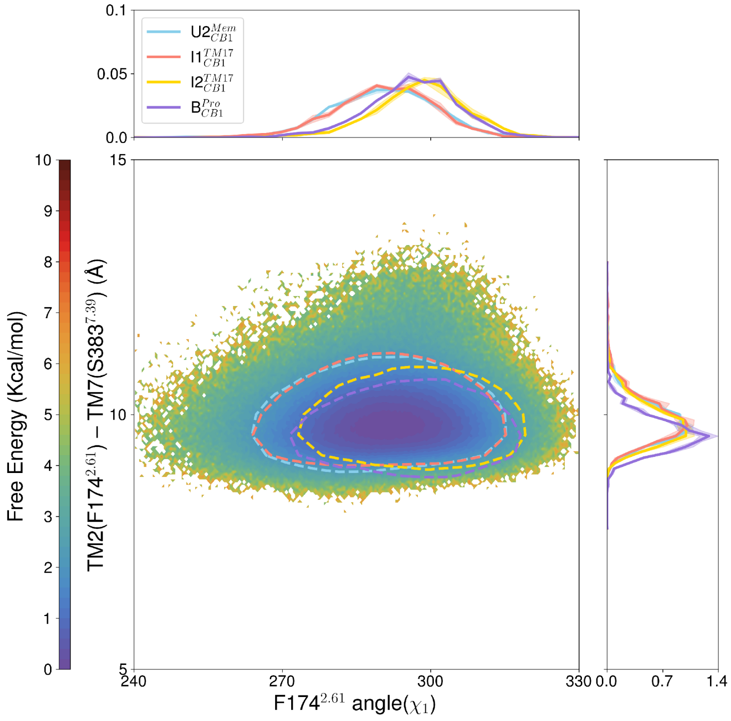


Figure S12: For $CB_1$, MSM weighted free energy landscape free energy landscape is plotted between TM2 - TM7 distance and F174$^{2.61}$ $\chi_1$ angle. Projections of important macrostate positions are shown as contour line on the landscape. Each feature of the free energy landscape is also plotted as probability density plot for different macrostates.

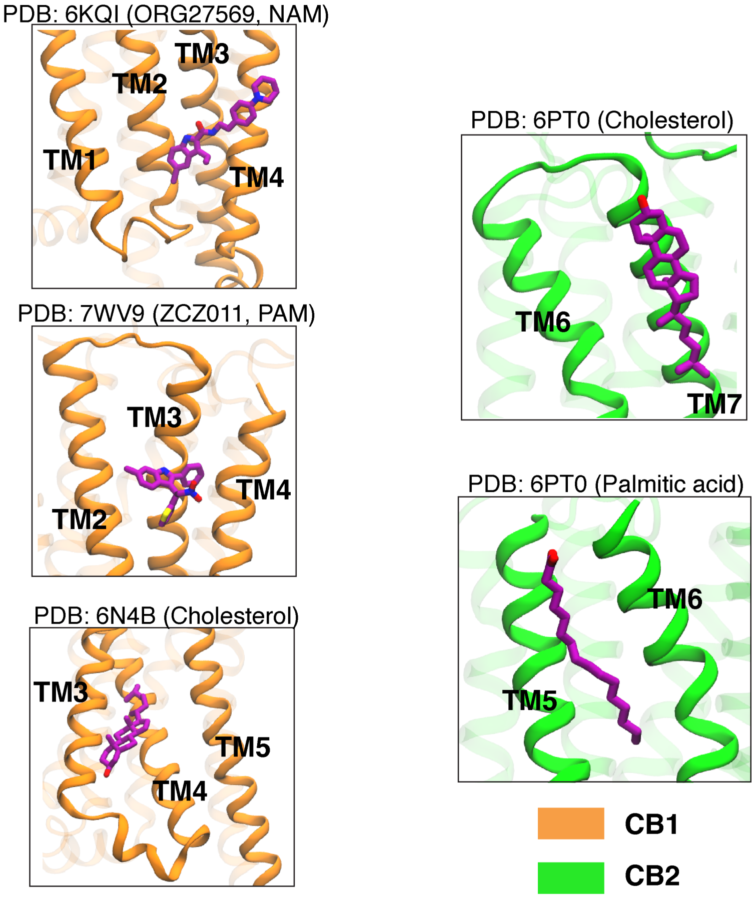


Figure S13: Allosteric sites found in the protein-lipid interface of the $CB_1$ (PDB: 6KQI,[2] 7WV9,[3] 6N4B[4]) (left panel) and $CB_2$ (PDB: 6PT0[5]) (right panel) crystal structures.

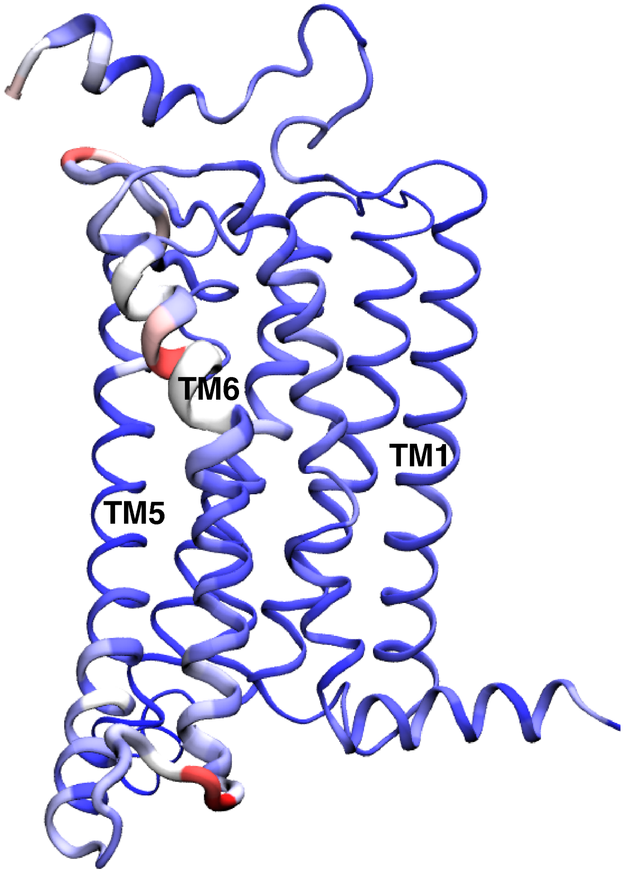


Figure S14: K-L divergences between $CB_2$ macrostate $U_{CB2}^{Mem}$ and $I1_{CB2}^{TM56}$ are shown as color and thickness gradients on the cartoon representation. Thickness gradients are shown as moving average.

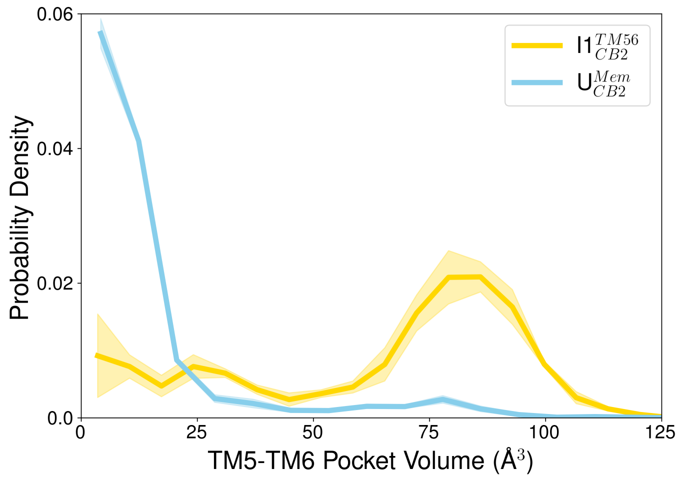


Figure S15: Probability density plot of pocket volume between TM5 and TM6 surface where the ligand binds inside the $CB_2$. Error in the probability density is obtained using bootstrapping method where each bootstrap sample consist of 1000 frames based on MSM weighted probability.

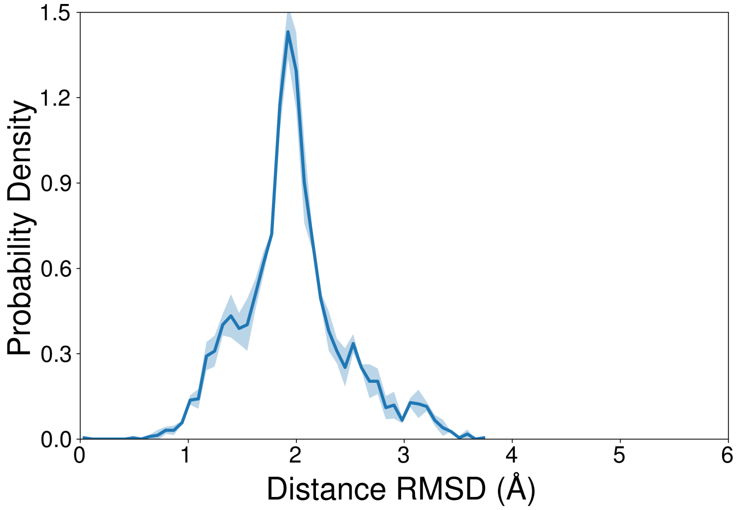


Figure S16: Probability density plot of distance RMSD for $CB_2$ macrostate $I2^{TM56}_{CB2}$. Error in distance RMSD calculations were calculated from on 3 bootstrap samples where each sample contains 1000 frames obtained from macrostate $I2^{TM56}_{CB2}$ based on MSM weighted probability.

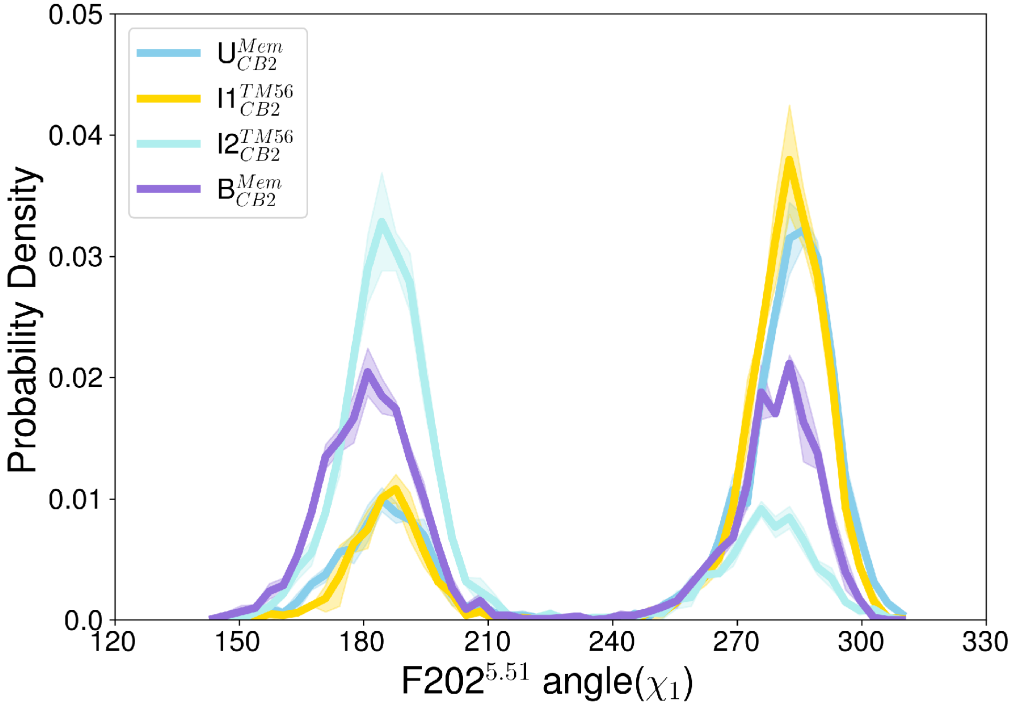


Figure S17: Probability density plot of $F202^{5.51}$ angle for different macrostates. Error in angle were calculated from on 3 bootstrap samples where each sample contains 1000 frames obtained from each macrostate based on MSM weighted probability.

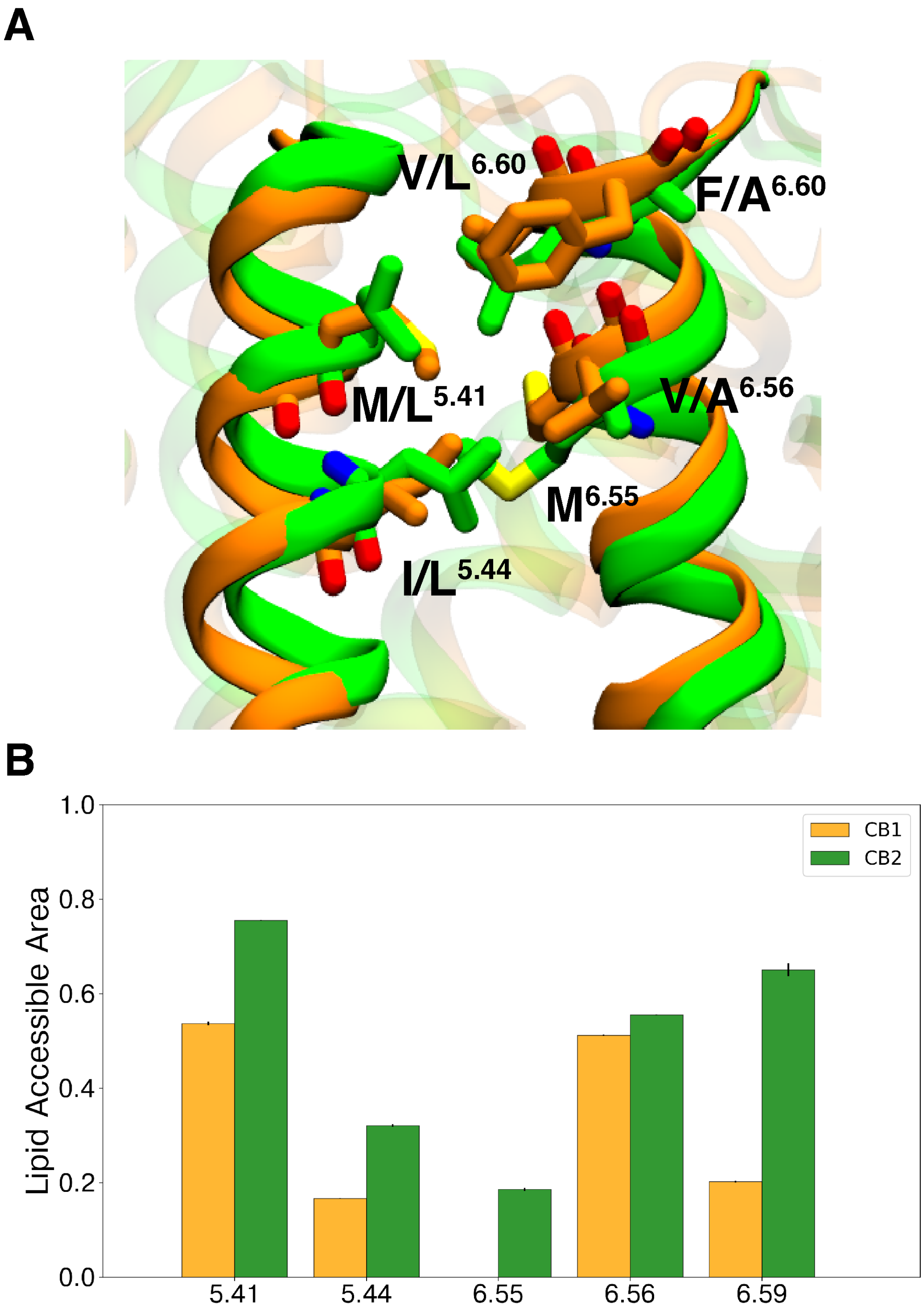


Figure S18: (A) Superposition of $CB_1$ (PDB: 5TGZ, color: orange) and $CB_2$ (PDB: 5ZTY, color: green) inactive structures to highlight the residue difference in the membrane surface of TM5 and TM6. (B) Surface area of the residue of interests are shown as bar plot for $CB_1$ (color: orange) and $CB_2$ (color: green).

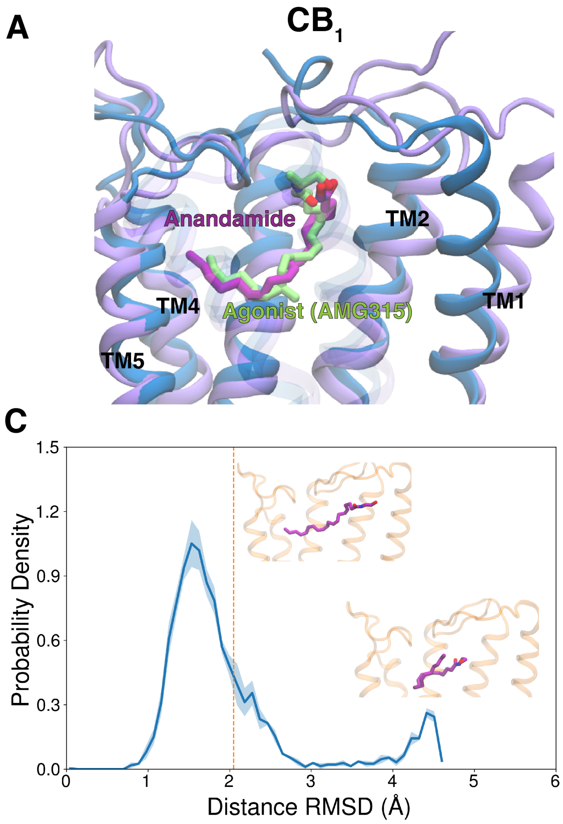


Figure S19: (A, B) Superposition of agonist (AMG315) bound active structure (PDB: 8GHV,[6] color: blue) and representative structure from $B_{CB1}^{Pro}$ (A) and $B_{CB2}^{Pro}$ (B) (color: mediumpurple). Ligands (Anandamide: purple; AMG315: lime) and proteins are shown as sticks and cartoon representations. Transparent representation of TM6 and TM7 are used to show the ligand binding position. (C, D) Probability density plot of distance RMSD for $B_{CB1}^{Pro}$ (C) and $B_{CB2}^{Pro}$ (D). Inset plots show the anandamide bound poses from the two peaks. Error in distance RMSD calculations were calculated from on 3 bootstrap samples where each sample contains 1000 frames obtained from macrostate B based on MSM weighted probability. Distance RMSD of anandamide which have similar pose as cryo-EM structure of AMG315 are shown as vertical dotted line.

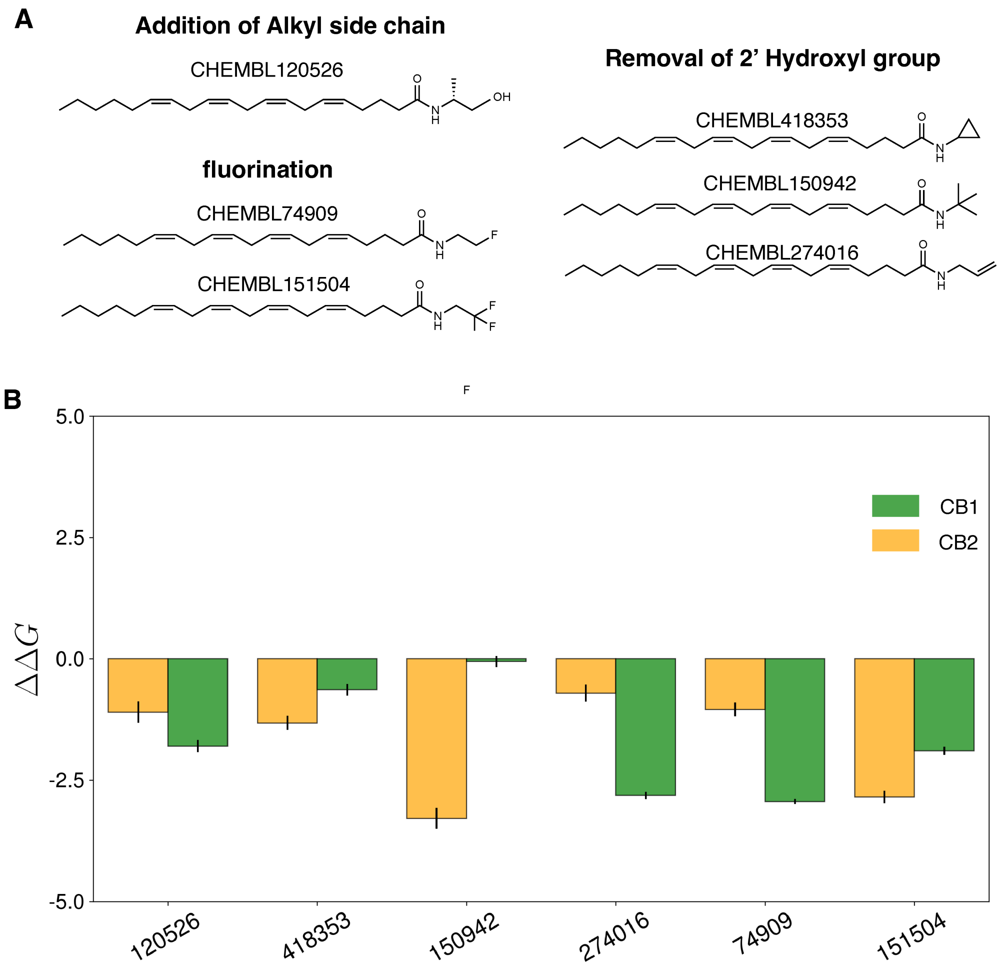


Figure S20: (A) Structures of different categories of anandamide analogs are shown with their CHEMBL database numbers. (B) Bar plots show the ΔΔG (kcal/mol) for $CB_1$ (color: orange) and $CB_2$ (color: green) for the ligands shown in panel (A).

**A**

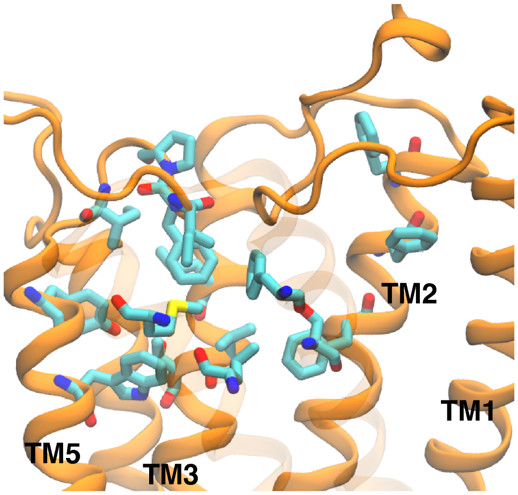


**Binding Pocket Residues**

| TM2 | TM3 | ECL2 | TM5 | TM6 | TM7 |
|---|---|---|---|---|---|
| F170$^{2.57}$ | L193$^{3.29}$ | I267$^{ECL2}$ | Y275$^{5.39}$ | L359$^{6.51}$ | F379$^{7.35}$ |
| F174$^{2.61}$ | V196$^{3.32}$ | F268$^{ECL2}$ | W279$^{5.43}$ | M363$^{6.55}$ | S383$^{7.39}$ |
| F177$^{2.64}$ | T197$^{3.33}$ | P269$^{ECL2}$ | | | |
| | | I271$^{ECL2}$ | | | |

**B**

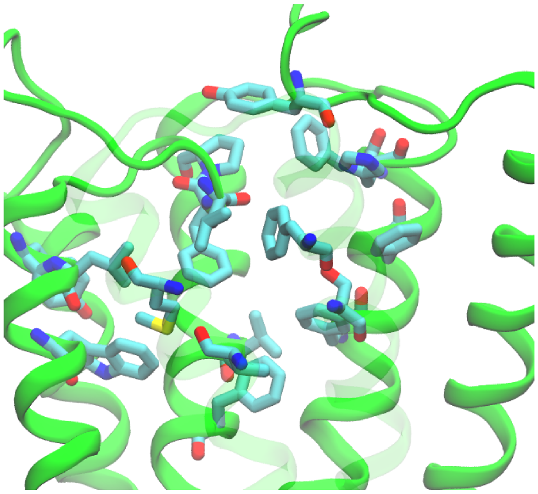

**Binding Pocket Residues**

| N-term | TM2 | TM3 | ECL2 | TM5 | TM6 |
|---|---|---|---|---|---|
| Y25$^{N\text{-}term}$ | F87$^{2.57}$ | V113$^{3.32}$ | L182$^{ECL2}$ | Y190$^{5.39}$ | V261$^{6.51}$ |
| | F91$^{2.61}$ | F117$^{3.36}$ | F183$^{ECL2}$ | L191$^{5.40}$ | M265$^{6.55}$ |
| | F94$^{2.64}$ | | P184$^{ECL2}$ | W194$^{5.43}$ | |
| | H95$^{2.65}$ | | | | |

| TM7 |
|---|
| F281$^{7.35}$ |
| S285$^{7.39}$ |

Figure S21: $CB_1$ (A) and $CB_2$ (B) structures with binding pocket residues that are used for ligand distance calculations. Protein structures are shown as cartoon and binding pocket residues are represented as sticks. TM6 and TM7 are represented as transparent.

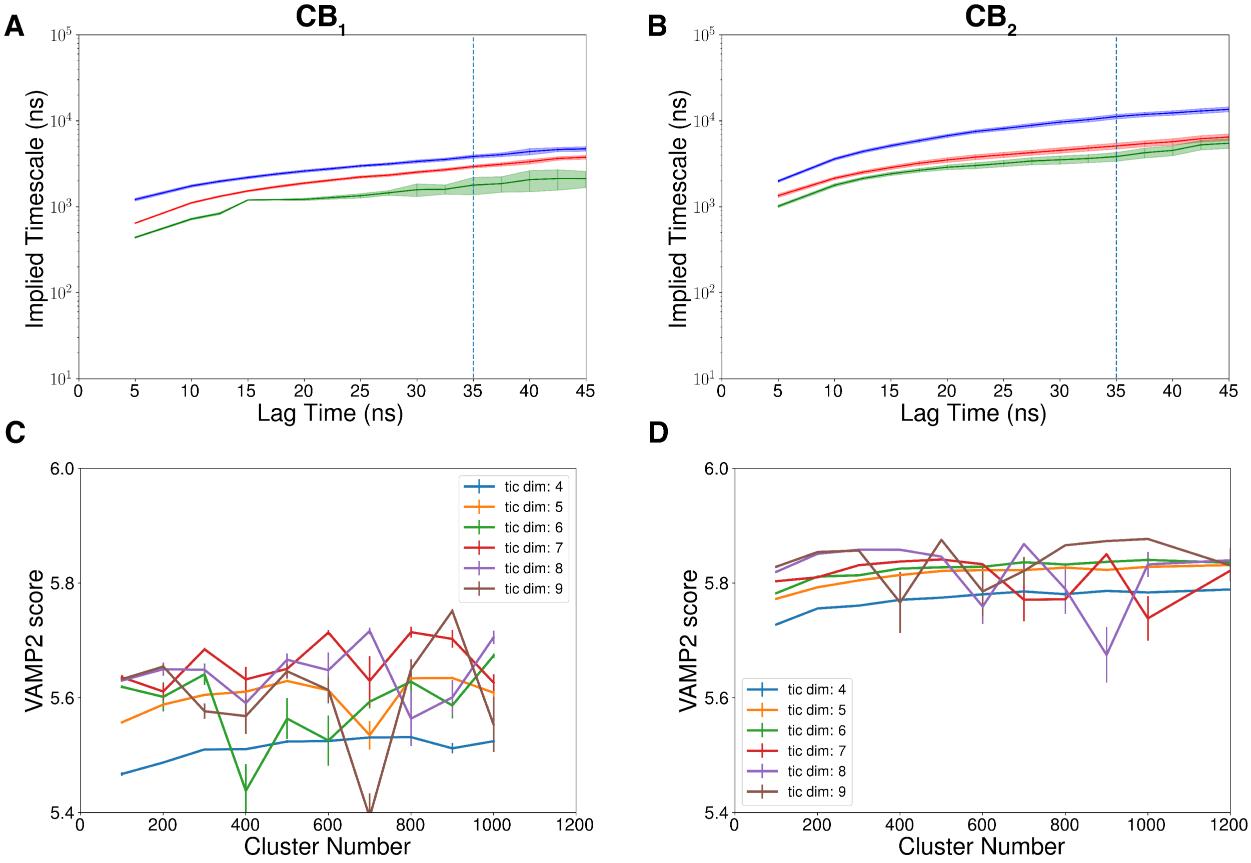


Figure S22: Implied timescales of the MSM with optimized hyperparameter for $CB_1$ (A) and $CB_2$ (B) are plotted against lagtime. Three slowest processes are shown are shown in this plot. Final lagtime for both systems is shown as a dotted line. VAMP-2 scores are plotted against number of states considered for $CB_1$ (C) and $CB_2$ (D) MSM building. VAMP-2 score variations based on the different tIC components are shown in different colors. For $CB_1$, optimized MSM has 800 states and 7 tICs. For $CB_2$, optimized MSM has 1000 states and 6 tICs.

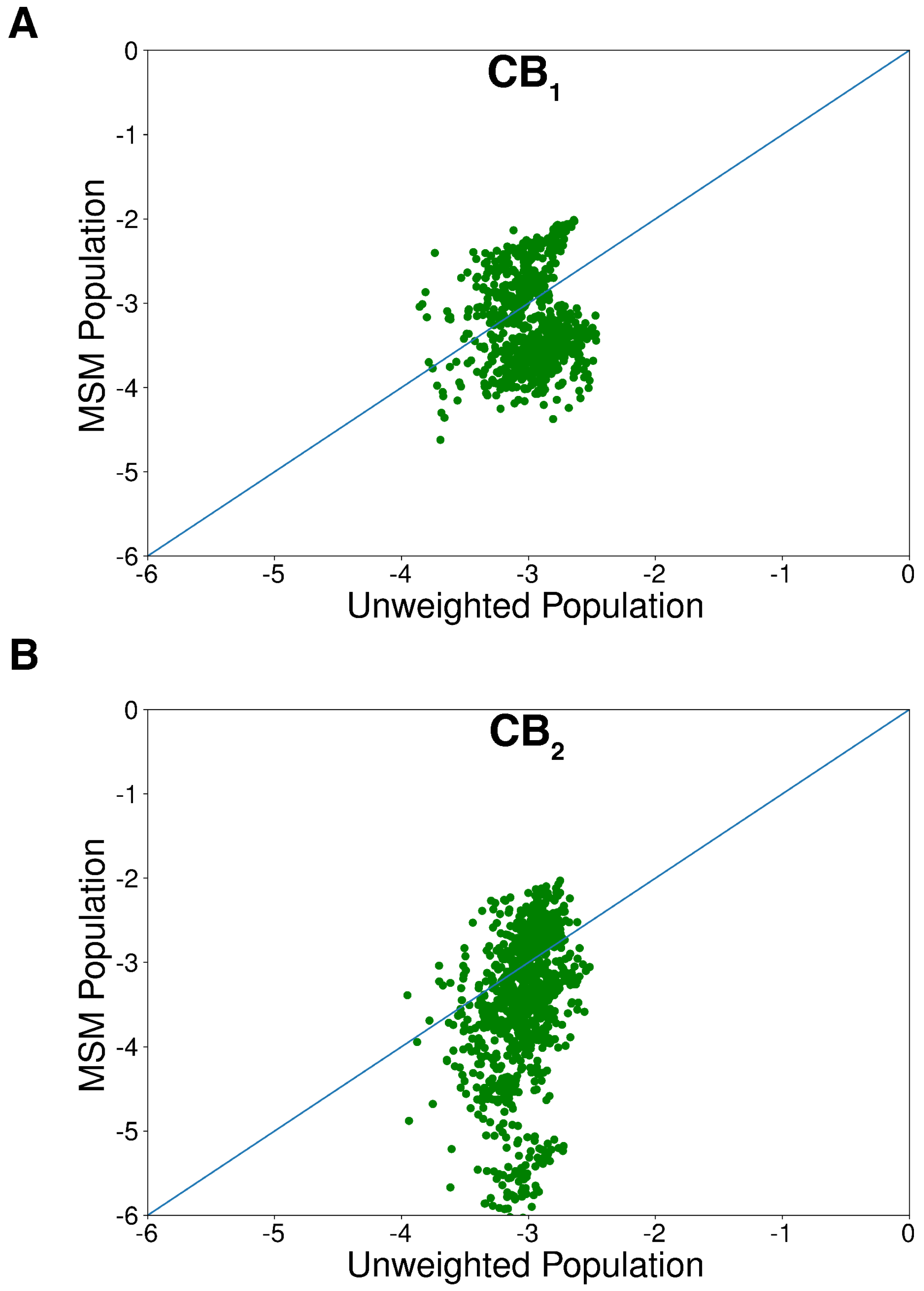


Figure S23: MSM weighted population are plotted against the raw population of the Markovian states for $CB_1$ (A) and $CB_2$ (B)

Table S1: Assymetric average brain membrane composition used in MD simulation for $CB_1$

| Head Group | Lipid Group | Upper Leaflet | Lower Leaflet | Total |
|---|---|---|---|---|
| phosphatidylcholine | DPPC | 8 | 5 | 13 |
| | POPC | 14 | 7 | 21 |
| | DOPC | 4 | 2 | 6 |
| | PAPC | 7 | 4 | 11 |
| | PDoPC | 1 | 0 | 1 |
| phosphatidylethanolamine | POPE | 2 | 4 | 6 |
| | PAPE | 5 | 9 | 14 |
| | PDoPE | 8 | 15 | 23 |
| sphingolipid | SSM | 9 | 3 | 12 |
| | OSM | 1 | 0 | 1 |
| | NSM | 2 | 0 | 2 |
| phosphatidylserine | DPPS | 0 | 1 | 1 |
| | POPS | 0 | 6 | 6 |
| | PAPS | 0 | 6 | 6 |
| Glycolipid | GM1 | 2 | 0 | 2 |
| | GM3 | 2 | 0 | 2 |
| phosphatidylinositol | POPI | 0 | 5 | 5 |
| | PIPI | 0 | 2 | 2 |
| Ceramide | CER180 | 1 | 1 | 2 |
| Sterol | Cholesterol | 62 | 58 | 120 |
| Total | | 128 | 128 | 256 |

Table S2: Assymetric average brain membrane composition used in MD simulation for $CB_2$

| Head Group | Lipid Group | Upper Leaflet | Lower Leaflet | Total |
|---|---|---|---|---|
| phosphatidylcholine | POPC | 16 | 8 | 24 |
| | DOPC | 1 | 1 | 2 |
| | PEPC | 1 | 0 | 1 |
| | PAPC | 4 | 2 | 6 |
| | PLPC | 23 | 11 | 34 |
| | PDoPC | 1 | 0 | 1 |
| | DAPC | 1 | 1 | 2 |
| phosphatidylethanolamine | POPE | 2 | 8 | 10 |
| | DOPE | 1 | 3 | 6 |
| | PAPE | 2 | 8 | 10 |
| | DAPE | 1 | 5 | 6 |
| | PLPE | 1 | 6 | 7 |
| | PDoPE | 1 | 3 | 4 |
| | DDoPE | 0 | 1 | 1 |
| sphingolipid | SSM | 15 | 7 | 22 |
| | OSM | 1 | 0 | 1 |
| | NSM | 9 | 5 | 14 |
| phosphatidylserine | POPS | 0 | 4 | 4 |
| | PAPS | 0 | 8 | 8 |
| | PLPS | 0 | 1 | 1 |
| phosphatidylinositol | POPI | 0 | 3 | 3 |
| | PIPI | 0 | 3 | 3 |
| phosphatidic acid | POPA | 0 | 1 | 1 |
| | PAPA | 0 | 1 | 1 |
| | PIPA | 0 | 1 | 1 |
| Ceramide | CER180 | 1 | 0 | 1 |
| Glycolipid | GM1 | 3 | 0 | 3 |
| | GM3 | 3 | 0 | 3 |
| Sterol | Cholesterol | 41 | 37 | 78 |
| diacylglycerol | POGL | 1 | 0 | 1 |
| Total | | 129 | 128 | 257 |
